# Transposable element abundance subtly contributes to lower fitness in maize

**DOI:** 10.1101/2023.09.18.557618

**Authors:** Michelle C. Stitzer, Merritt B. Khaipho-Burch, Asher I. Hudson, Baoxing Song, Jose Arcadio Valdez-Franco, Guillaume Ramstein, Cedric Feschotte, Edward S. Buckler

## Abstract

Transposable elements (TEs) have long been shown to have deleterious effects on the survival and reproduction of their host organism. As TEs are mobile DNA that jump to new positions, this deleterious cost can occur directly, by inserting into genes and regulatory sequences. Classical population genetic theory suggests copy-number dependent selection against TEs is necessary to prevent TEs from expanding so much they take over a genome. Such models have been difficult to interpret when applied to large genomes like maize, where there are hundreds of thousands of TE insertions that collectively make up 85% of the genome. Here, we use nearly 5000 inbred lines from maize mapping populations and a pan-genomic imputation approach to measure TE content. Segregating TE content gives rise to 100 Mb differences between individuals, and populations often show transgressive segregation in TE content. We use replicated phenotypes measured in hybrids across numerous years and environments to empirically measure the fitness costs of TEs. For an annual plant like maize, grain yield is not only a key agronomic phenotype, but also a direct measure of reproductive output. We find weak negative effects of TE accumulation on grain yield, nearing the limit of the efficacy of natural selection in maize. This results in a loss of one kernel (≈0.1% of average per-plant yield) for every additional 14 Mb of TE content. This deleterious load is enriched in TEs within 1 kilobase of genes and young TE insertions. Together, we provide rare empirical measurements of the fitness costs of TEs, and suggest that the TEs we see today in the genome have been filtered by selection against their deleterious consequences on maize fitness.

## Introduction

Across eukaryotes, the amount of nuclear DNA varies by five orders of magnitude. These differences do not scale with eukaryotic organismal complexity, and whether differences in genome size are adaptive or shaped by genetic drift and evolving neutrally remain contentious (Elliott and Gregory, 2015). The underlying cause of these interspecific differences in genome size lies in the amount of nongenic DNA, almost certainly a consequence of past transposable element (TE) activity. When TEs jump to a new position, they generate an insertion of their own DNA sequence, and uncontrolled transposition can generate extreme costs to host genomes when TE insertions in genes and other functional sequences disrupt cellular function. TEs are thus selfish DNA and parasitic to their host genome (Orgel and Crick, 1980; Doolittle and Sapienza, 1980).

The maize genome contains over 350,000 TEs (Stitzer *et al*., 2021), and fragments derived from TEs cumulatively make up over 85% of the genome (Schnable *et al*., 2009; Hufford *et al*., 2021). These TEs come from over 27,000 families in all known plant TE superfamilies, and are found in variable genic, chromatin, methylation, and recombinational environments within the genome (Baucom *et al*., 2009; Stitzer *et al*., 2021). Maize TEs are extremely polymorphic, with only half of TE insertions shared at the same position between any two individuals (Brunner *et al*., 2005; Morgante *et al*., 2005; Anderson *et al*., 2019; Munasinghe *et al*., 2023). The high abundance and diversity of TEs in maize, as compared to previously investigated model taxa, suggests the fitness costs of TEs cannot possibly be as high as those previously measured in yeast and flies. Although maize is often cited as an example of a large genome, it is in fact below the average genome size of both plants (6.1 Gb) and animals (4.2 Gb) (Dodsworth *et al*., 2015; Elliott and Gregory, 2015). Quantifying the fitness costs of maize TEs enriches our understanding of the forces acting on TEs in a more typical eukaryotic genome.

Here, we combine measurements of TE content in maize mapping populations with fitness measurements of these individuals, to reveal that there is weak but pervasive negative selection acting on TEs in maize. Selection against TEs lies at the boundary between drift and selection in maize, suggesting that although the maize genome is relatively large, its size is likely constrained by the cumulative deleterious effects of the pesky TEs that make up the bulk of its sequence.

## Results and Discussion

Genome size is determined by the combination of DNA sequences inherited from both parents, in the absence of novel mutations. Transposition events are infrequent in maize pedigrees, where few to no insertions occur per generation in most germplasm (Dooner *et al*., 2019). Thus, genome size and TE content can be well-approximated by imputation from assembled parental genomes to genotyped progeny. We used a maize pan-genome (Bradbury *et al*., 2021; Valdes Franco *et al*., 2020) to impute genome size and TE content, by projecting TE annotations and haplotype blocks from parental genome assemblies to genotyped RILs. We did so in the US Nested Association Mapping (NAM) population of maize (McMullen *et al*., 2009; Gage *et al*., 2020), a set of 4,975 genotyped recombinant inbred lines (RILs) generated from twenty-five parental inbred lines crossed to the original reference genotype of maize (B73) then self-pollinated for 4-6 generations (McMullen *et al*., 2009). Many maize inbred lines were publicly released while still containing residual heterozygosity, and these regions subsequently became homozygous for alternative alleles in different lineages (McMullen *et al*., 2009; Liang and Schnable, 2016). We extensively filtered such regions of the genome that did not return the parents expected from the pedigree. While this filtering reduces total genome size, our imputed genome size is still highly correlated to flow cytometry metrics of DNA content (Supp Fig. S2). In RIL individuals, genome size varies by 97.8 megabases (Mb) (Fig 1A, Supp. Fig. S2), approximately 4.2 percent of the B73 reference genome size. Both genome size (Fig 1A) and TE content (Fig 1B) observed in the RILs vary by NAM biparental family, as each NAM parent contributes different alleles to the RIL progeny. We see transgressive segregation for both genome size and TE content in certain NAM families, such as CML322, where the majority of RILs have higher values than either parent (Fig 1A-B). This is likely contributed by high variance between parental genomes in alternative haplotype content. Much of the variation in genome size can be explained by the strong, positive relationship of genome size with TE content (Spearman’s correlation, rho=0.961, p<2.2e-16) (Fig 1C). Further, a high proportion (0.937, 95% CI 0.904-0.965) of variance in TE content can be explained by a kinship matrix describing relatedness of these RILs.

**Fig. 1.**
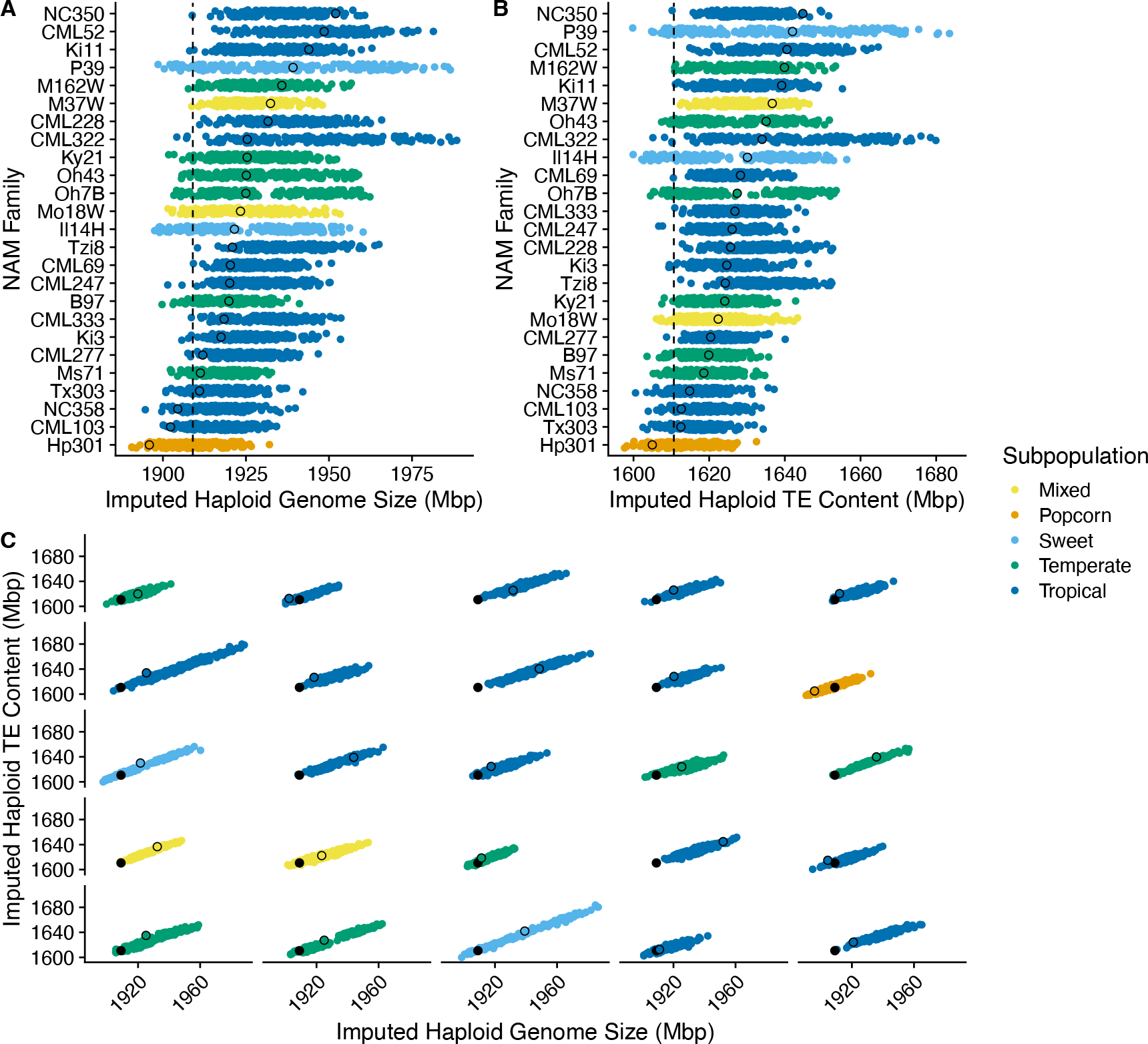
Imputed genome size and TE content for each NAM RIL. (A) Imputed haploid genome size for each NAM RIL, split by NAM family, (B) Imputed haploid TE content for each NAM RIL, split by NAM family (C) Relationship between imputed haploid genome content and imputed haploid TE content in each NAM RIL. Each colored point reflects a NAM RIL, colored by maize subpopulation. Black outlined circle is the parental value calculated from its genome assembly, and black filled circles and dashed line are values for the genome assembly of the B73 common parent.

To test whether genome size and TE content are associated with fitness related phenotypes, we use data from multi-environment yield trials of almost one million individual plants. These consist of F1 hybrids of each RIL with the former commercial tester, inbred line PHZ51 (Larsson *et al*., 2017; Ramstein *et al*., 2020). As modern corn breeding has focused on selection for alleles that perform well in a hybrid background, these phenotypes provide relevant effect estimates, particularly on traits related to fitness. We use phenotypes of flowering time and grain yield to measure fitness. For annual plants like maize and its wild relative teosinte, flowering time contributes to fitness, as the plant must appropriately incorporate environmental signals to flower synchronously with others, and before the end of the growing season. Fitness can more directly be measured as the number of viable seeds produced per plant, which is well-measured by grain yield. Maize breeding has targeted increased grain yield as measured through seed mass per field area (Hallauer *et al*., 1988; Duvick *et al*., 2003). To better assess genome content of these hybrids, we assembled the genome of the hybrid tester PHZ51, and added its genomic contribution to each RIL hybrid to generate a diploid value that reflects the diploid genome of the phenotyped F1 individuals. For all analyses, we use best linear unbiased estimator (BLUE) values for flowering time and grain yield calculated in Ramstein *et al*. (2020) for a subset of 1,723 RIL hybrids phenotyped in six environmental trials (Larsson *et al*., 2017). The mean TE content in the subset of phenotyped RILs is approximately 1 Mb larger than the 3,252 NAM RILs not phenotyped (difference = 1.24e+06, p<0.001; Supp. Fig S1). The phenotyped subset of RILs were originally chosen for similar flowering times to limit the effect of growing season length on grain yield, and in this experiment BLUE estimates of female flowering time (Days To Silking, DTS) range from 64 to 77.8 days (*µ*= 70.355, SD= 2.06). As previously shown (Ramstein *et al*., 2020), grain yield is negatively correlated with DTS (Supp. Fig. S3A), thus we introduce flowering time as a fixed effect in the calculation of grain yield BLUEs, to correct for flowering time (Supp. Fig. S3). These adjusted grain yield (GY) values range from 1.866 to 10.415 tonnes/hectare (*µ*= 6.99, SD= 0.86).

We first associate genome size to flowering time, as multiple reports have shown that maize individuals with larger genomes flower later (Rayburn *et al*., 1994; Jian *et al*., 2017; Bilinski *et al*., 2018; Li *et al*., 2018). Consistent with previous work, we find that RIL hybrids with larger genomes flower later (Figure 2A). This association has previously been proposed as an effect of simply having more DNA that takes time to replicate (Bilinski *et al*., 2018), or specific effects from repeat classes like chromosomal knobs (Jian *et al*., 2017), ribosomal repeats (Li *et al*., 2018), or telomeres (Choi *et al*., 2021). To test the impact of components of genome size, we fit a series of linear models predicting female flowering time (DTS) in these hybrids from different repetitive categories (Table 1). These include the amount of TE base pairs in their genome, the amount of four tandem repeat class base pairs (ribosomal, centromeric, telomeric, and knob), and amount of the genome coming from the B73 parent (Table 1). The model explains a statistically significant and substantial proportion of variance (R2=0.133, p < 2.2e-16) and all significant positive associations of repeat classes with flowering time are positive, except for TE base pairs which is negative (Table 1). We additionally fit a model using three principal component (PC) terms from a kinship matrix to correct for population structure, which recovers similar effects (Supp. Table S1). Much of the early evolution of maize occurred in tropical environments, but dispersal to temperate environments over the last several millennia required major flowering time and photoperiod adaptations (Swarts *et al*., 2017). Additionally, germplasm from tropical environments tends to have larger genomes (Chia *et al*., 2012; Hufford *et al*., 2021). To detect whether the relationship varies across different biparental families, we fit models within each NAM family. For flowering time, 58% (14) of NAM families show positive effects of TE base pairs on DTS, (Figure 3A), and positive effects combined using Fisher’s method are significant (p = 0.014). Although we see an overall negative effect of TE base pairs when considering all NAM families, within individual NAM families we see more balanced effects on flowering time. In other taxa (e.g. wheat, *Capsella, Arabidopsis, Brassica*; Yan *et al*., 2006; Nitcher *et al*., 2014; Niu *et al*., 2019; Baduel *et al*., 2018; Quadrana, 2020; Cai *et al*., 2022) and within maize (Salvi *et al*., 2007; Hung *et al*., 2012; Yang *et al*., 2013; Huang *et al*., 2018), large effect TE insertions consistently accelerate flowering by disrupting regulation of key flowering time pathway genes. Our results are in line with a polygenic view of such disruption of flowering, arising from quantitative variation in TE content with individually small effects on flowering time. Notably, the regression coefficient for TE bp implies 44 Mb of additional TE content would accelerate flowering only by one day – an effect size similar to the largest single locus QTLs segregating in this population (Buckler *et al*., 2009). Additionally, the presence of active TEs in maize has been associated with earlier flowering, hypothesized to be due to activation of general stress pathways (Skibbe *et al*., 2009). In total, TEs seem to have a disproportionate impact on flowering time beyond simply being made of DNA.

**Table 1.** Relationship between genomic repeats and flowering time (DTS) and grain yield (GY).

|  | DTS | GY |
| --- | --- | --- |
| (Intercept) | $6.64 \times 10^{1***}$ | $2.57 \times 10^{1**}$ |
| TE bp | $-2.27 \times 10^{-8***}$ | $-6.93 \times 10^{-9*}$ |
| nonTE, nonRepeat bp | $1.48 \times 10^{-7***}$ | $8.75 \times 10^{-9}$ |
| Knob bp | $1.62 \times 10^{-7***}$ | $1.80 \times 10^{-8*}$ |
| Centromere bp | $3.64 \times 10^{-7***}$ | $-2.99 \times 10^{-8}$ |
| Telomere bp | $6.29 \times 10^{-7*}$ | $2.09 \times 10^{-7}$ |
| Ribosomal bp | $6.90 \times 10^{-7*}$ | $4.49 \times 10^{-7**}$ |
| B73 bp | $-1.89 \times 10^{-9***}$ | $-1.33 \times 10^{-10}$ |
| Num.Obs. | 1561 | 1454 |
| R2 | 0.133 | 0.018 |
| R2 Adj. | 0.129 | 0.013 |

+ p < 0.1, \* p < 0.05, \*\* p < 0.01, \*\*\* p < 0.001

**Fig. 2.**
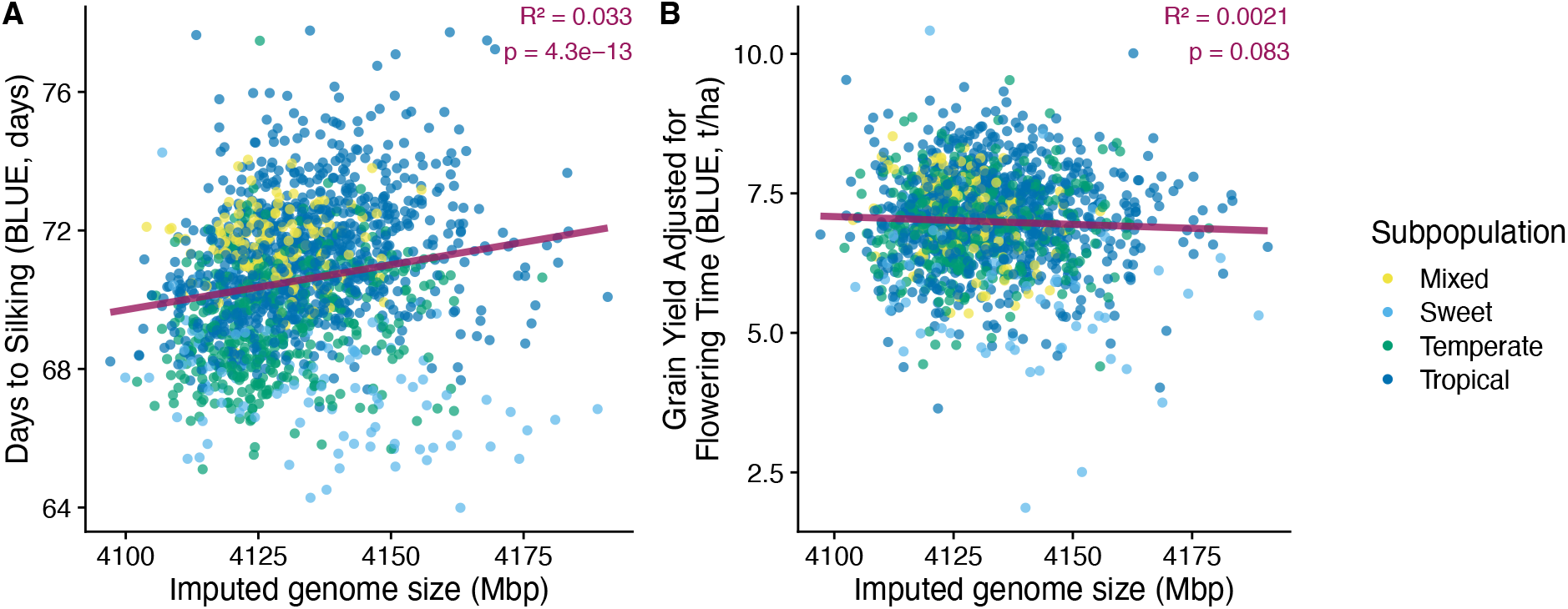
Phenotype associations with imputed genome size. (A) Imputed diploid (2N) hybrid genome size vs Days to Silking, (B) Imputed diploid (2N) hybrid genome size vs Grain Yield corrected for flowering time. Each colored point reflects a NAM RIL, colored by maize subpopulation. Lines show linear regression.

**Fig. 3.**
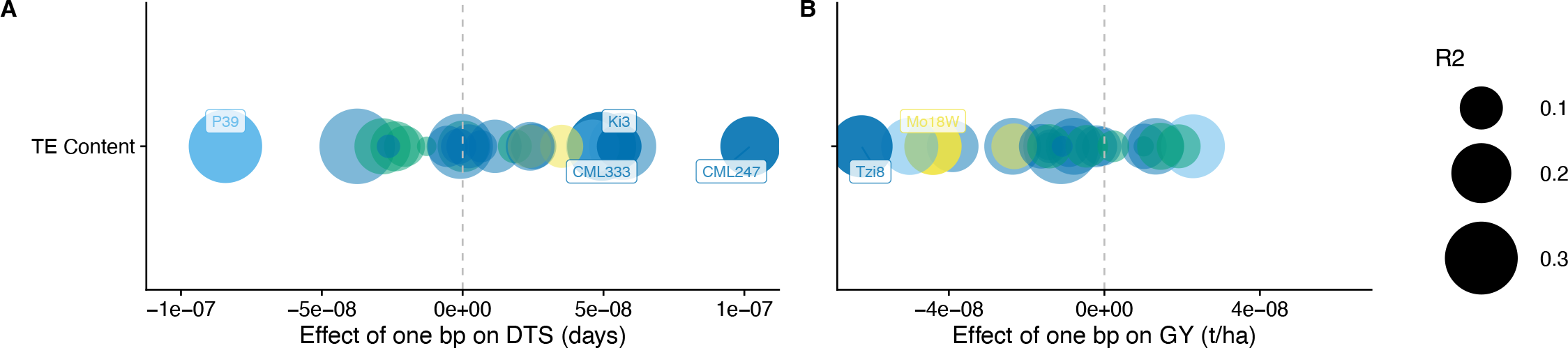
Phenotype associations with transposable element content, for each NAM family. (A) Female flowering time (Days to Silking), (B) Grain yield. Each colored point reflects the regression coefficient of a NAM family, colored by maize subpopulation, and point size reflects the coefficient of determination (R2) for the entire model. Families that are significantly associated are plotted in a darker color, and labeled with their family name. All model coefficients are plotted in Supp. Figure S4, and model summaries in Supp. Tables S3 and S4.

Although flowering is essential to fitness, grain yield more directly quantifies maize fitness, and has been a target of selection for millennia. As expected for fitness, the genetic architecture underlying grain yield is highly polygenic, with few validated yield QTL segregating in maize breeding populations (Giraud *et al*., 2017; Ramstein *et al*., 2020; Simmons *et al*., 2021; Khaipho-Burch *et al*., 2023). Unlike flowering time, little is known about the relationship between grain yield and genome size. We see a non-significant negative relationship, where individuals with larger genomes show lower fitness (Figure 2B). As with flowering time, we fit linear models predicting grain yield (GY) from TE and various types of non-TE repetitive DNA, controlling for population structure with the amount of the genome coming from the common B73 parent. The model explains a statistically significant but weak amount of variation in grain yield (R2=0.018, p=0.0004; Table 1). A similar model using PCs of a kinship matrix for population structure control explains more variance, but effect sizes remain similar (R2=0.068, p < 2.2e-16; Supp Table S1). In both models, higher TE content is significantly associated with reduced grain yield, while higher abundance of ribosomal and knob repeats are significantly associated with increased grain yield only without PC correction for population structure (Table 1). Quicker protein production with more abundant ribosomes may accelerate growth rates, although it is not clear how much compensation occurs between genomic rDNA copy number and rRNA expression (Li *et al*., 2018). Similarly, a fitness benefit of chromosomal knobs has been previously implicated in models explaining equilibrium frequencies of meiotic drive in maize (Hall and Dawe, 2018). The negative association of TE abundance with fitness confirms theoretical and empirical evidence of the deleterious costs of TEs (Charlesworth and Charlesworth, 1983; Charlesworth and Langley, 1989). We replicate this negative association between TE abundance and grain yield within individual NAM families, and find a negative effect for the majority of NAM families. Individuals with more TEs show lower grain yield, in 70% (17) of NAM families (Figure 3B), and a combined p-value using Fisher’s method shows a significant negative association between TE base pairs and fitness (p = 0.033). Making a number of simplifying assumptions about average per-plant yield (see Methods), our models predict a loss of one kernel’s worth of yield for every additional 14.43 Mb of TEs. Making further assumptions about the average length of a TE, this fitness cost reflects a selection coefficient against individual TE insertion of s=-1.4e-7. The per-locus impact of TEs on fitness in maize is much smaller than seen in other taxa. Quite simply, the maize genome could not exist if every one of the 350,000 TEs were reducing fitness by 0.1% to 5%, as seen in fly and yeast experiments (Wilke and Adams, 1992; Mackay, 1989; Pasyukova *et al*., 2004). The proximate cause of these small differences in fitness are small enough to be due to the bioenergetic cost to replicate the additional TE DNA. As such, differences between taxa in genome size, TE content, and population history likely affect how selection acts on TEs. The shift between regimes where genetic drift dictates the fate of an allele and where selection is effective lies at 2*N s*, where *N* is the effective population size, and *s* is the selection coefficient (Kimura, 1962). Maize underwent a population bottleneck as it was domesticated from its wild relative teosinte, and the effective population size is estimated around 100,000 (Beissinger *et al*., 2016). Our crude estimate of *s* thus suggests that 2*Ns* = 0.28, well within the range of estimated values where genetic drift will predominate, and selection cannot push TE content even lower.

To better understand the properties of TEs that may underlie these effects, we consider a number of characteristics previously shown to impact how deleterious a TE is to host fitness. We present the relationship with grain yield in the text, as it best explains fitness, although associations with flowering time are presented in Supplemental Tables S5, S6, and S7. One relationship that has been widely supported is that TEs close to genes may disrupt host fitness to a greater effect than those far from genes (Medstrand *et al*., 2002; Wright *et al*., 2003; Hollister and Gaut, 2009). We partition TE bp to those TEs inside of a gene model, 1 kb from a gene, 1 - 5 kb from a gene, and all other TEs greater than 5 kb from a gene, then fit a linear model relating TE bp in these categories to grain yield. This model explains more variance (R2=0.040, p=1.941e-11) than previous models considering TEs in bulk. While the majority of TE bp is found greater than 5 kb away from genes (mean 2,980.7 Mb diploid, 83.3% of all TE base pairs), this does not show a significant association with grain yield (Figure 4A, Supp. Table S5). There are significant positive effects for TE base pairs inside of genes, most often intronic insertions (mean 73.7 Mb, 2.1% of all TE bp). This positive effect is contrary to the expectation that TEs that insert within genes will have a high deleterious cost, as they are likely to disrupt transcription, splicing, or even coding sequence of that gene (Lisch, 2013; Hirsch and Springer, 2017; Wells and Feschotte, 2020). Our observation could reflect a filtering process, where new genic insertions are rapidly removed by selection, leaving only TEs in genes that have neutral or even conditionally beneficial fitness effects. Consistent with such a model, Qiu *et al*. (2021) find TE insertions within genes are found at elevated frequencies in a maize diversity panel compared to insertions elsewhere in the genome. A positive effect of TE bp on grain yield is also seen for TE bp one to five kb from a gene (mean 326.0 Mb, 9.1% of all TE bp). In maize, this region encompasses the transition from genic euchromatin to silenced heterochromatin (Li *et al*., 2015; Martin *et al*., 2021). For many genes, this occurs due to the presence of an ‘island’ of CHH methylation, which prevents genic euchromatin from spreading out to TE-dense regions (Li *et al*., 2015; Martin *et al*., 2021). Higher load of TEs in this region 1-5 kb from genes may thus strengthen the differentiation between compartments of the maize genome, enforcing silencing of TEs. The only subset of TEs significantly negatively associated with grain yield are those within 1 kb of a gene (mean 93.7 Mb, 2.6% of all TE bp). It has been extensively shown that TE insertions can alter gene expression (Lisch, 2013; Hirsch and Springer, 2017; Uzunovic’
s *et al*., 2019), either due to disruption of existing regulatory sequence, or the contribution of new regulatory sequence encoded by the TE. In total, these analyses partitioning TE bp into distance classes from genes suggests that the negative impact of TEs on fitness is predominantly due to TE bp in this close regulatory space near genes. It is important to note that although this group has the largest negative effect, its burden is smaller in magnitude than other classes as they only make up 2.6% of all TE base pairs.

**Fig. 4.**
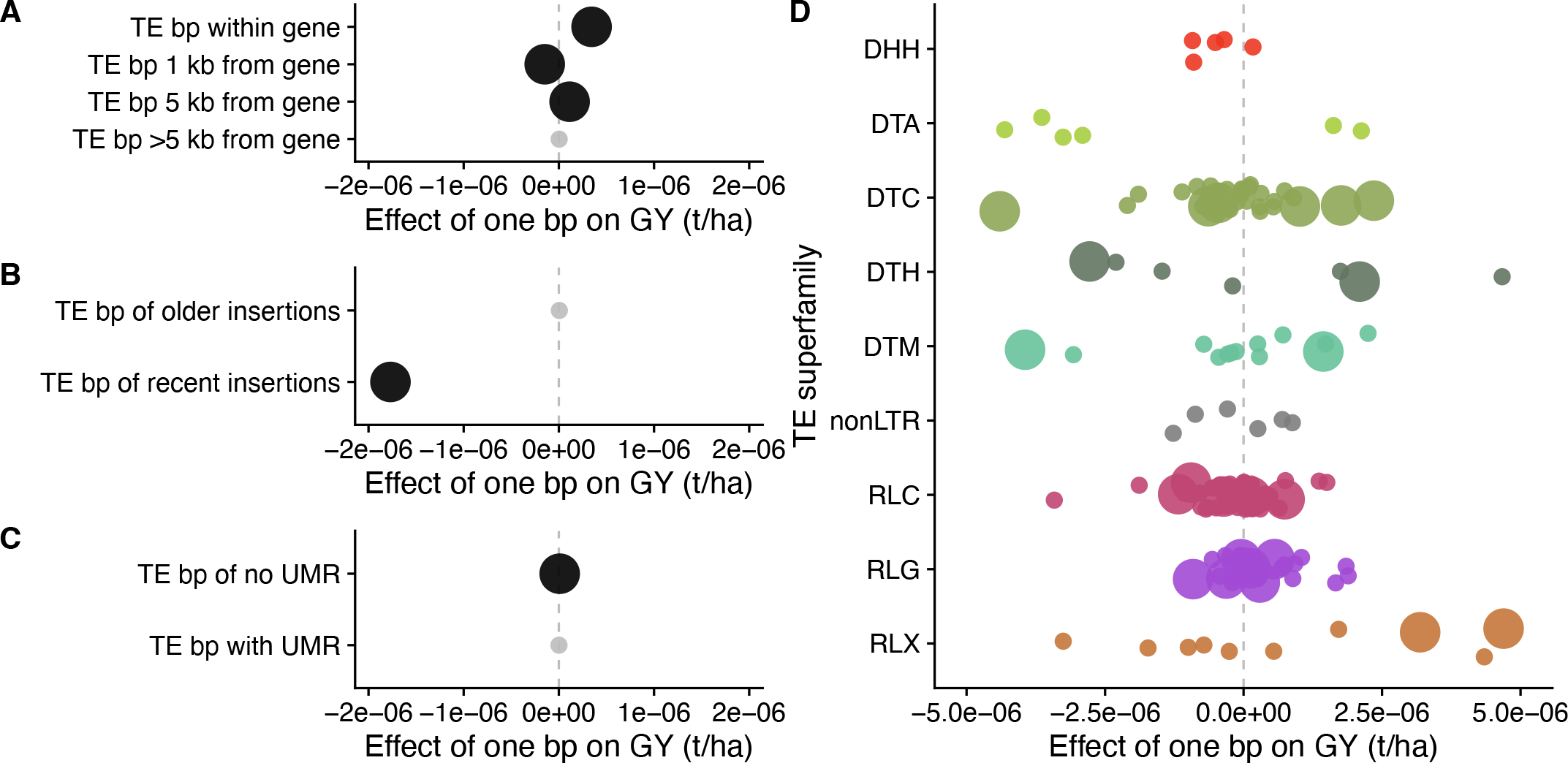
GY associations with different subsets of TE base pairs. Effect size of associations of TE base pairs with grain yield, for distance to gene (A), age of insertion (B), presence of unmethylated regions (UMR) (C), and TE families (D). Significant effects (p*<*0.05) shown with larger circle. Small gray points in A-C lack statistical significance, as do small points in D.

Actively jumping TEs generate mutations that selection may not yet have had a chance to remove from populations. To measure the fitness consequences of recent TE insertions, we measure which TEs have accumulated no substitutions from their consensus copy, and contrast this to older, degraded TEs with at least one substitution. We require insertions to be a minimum of 500 bp long, as this means these recent insertions are younger than 30,000 years, given a mutation rate of 3.3e-8 substitutions/generation (Clark *et al*., 2005). When we associate these two classes of TEs with grain yield, only recent insertions have a significant negative effect on yield (Figure 4B, Supp. Table S6), and the model explains similar variance to that of models containing all repeat classes (R2=0.016, p=3.536e-05). This suggests that old segregating TEs have minimal impact on yield, much less than that observed for recent insertions that are not yet purged from maize populations.

The deleterious cost of TEs near genes has been shown to be more extreme when TEs near genes are silenced by DNA methylation (Hollister and Gaut, 2009; Choi *et al*., 2021), but unmethylated TEs also pose a deleterious cost due to their ability to jump to new positions and make mutations. To test the effect of methylation state of TEs on fitness, we use measures of unmethylated regions (UMRs) of the maize NAM genome assemblies (Hufford *et al*., 2021). These unmethylated regions likely encompass accessible chromatin regions across maize tissues (Crisp *et al*., 2020). We find a significant positive effect of load of methylated TEs (those without a UMR), and a nonsignificant positive effect for those TEs that contain a UMR on grain yield (Figure 4C, Supp. Table S7). This model explains very little variance in grain yield, and does not reach statistical significance (R2=0.005, p=0.0638). While more subtle methylation differences between maize individuals surely exist, it is not clear whether methylation can be imputed to RILs, and future studies directly measuring methylation in hybrids could address this.

Finally, we consider the effects of different TE families on fitness. Often considered as a single category of masked DNA, TEs are extremely variable in their replication mechanisms and impacts on their host genomes (Lisch, 2013; Wells and Feschotte, 2020). Although superfamilies of TEs demonstrate general genomic patterns and can be related to our phenotypes (Supp Table S2), the true unit of selection for a TE is that of closely related lineages of TE families. Maize TEs vary in their genomic position, activity, and age, which is best captured at the level of TE family (Stitzer *et al*., 2021). To ensure we are capturing genome-wide signal for each TE family, and not simply linkage to flowering or grain yield QTL, we analyze the 170 TE families that account for greater than 10 Mb of DNA sequence across all NAM parent assemblies. These 170 TE families belong to 11 different superfamilies of TEs, that are representative of TE diversity in the maize genome (Stitzer *et al*., 2021). We simultaneously estimate the effect of each TE family on grain yield by fitting a linear model with terms summarizing the base pairs of the TE family in each RIL (Supp. Table S8), as well as terms for the summed base pairs of all other TEs families and base pairs contributed by the B73 parent. This model contains more parameters, but also explains more variance in grain yield than any other model we fit (R2=0.265, p<2.2e-16). We find that 23 TE families are significantly associated with grain yield; 12 are associated with lower grain yield, while 11 TE families are associated with higher grain yield (Fig 4D), but none of these associations surpass FDR correction. Families associated with negative effects include DTC00048, an En/Spm family originally named Doppia, that was discovered when active transposition generated chromosomal rearrangements and gene duplications at the *r1* locus (Walker *et al*., 1995; Bercury *et al*., 2001). Families associated with positive effects include RLG00017, a Ty3 LTR retrotransposon family originally named Dagaf, that confers salt responsiveness to nearby genes (Makarevitch *et al*., 2015). The great variability of associations between TE family abundance and fitness highlights the great variability of TE families, and the limitations of summarizing all TE content into a single category. We are unable to test individually the impact of smaller TE families, but in aggregate these show a significant negative effect on fitness (effect estimate = -9.05e09; Supp. Table S8). Many of the actively transposing TEs in maize, particularly for LTR retrotransposons, come from families with a handful of copies in the genome (Dooner *et al*., 2019; Wessler, 2019). It is likely that these families may have disproportionately large effects on fitness, relative to their abundance, and future studies of TE deleterious load should develop strategies to test the effects of these rare variants.

## Conclusion

The mutagenic impact of *de novo* TE insertion has been extensively documented through genetic analysis of spontaneous mutants of many model species, including those in maize that led to the discovery of transposition itself (McClintock, 1950). While maize has an abundance of TEs, making up over 85% of the genome (Schnable *et al*., 2009; Hufford *et al*., 2021), TE content is extremely variable between maize individuals (Brunner *et al*., 2005; Morgante *et al*., 2005; Anderson *et al*., 2019; Munasinghe *et al*., 2023). However, most TEs present in the maize genome have not transposed recently – the median age of a TE insertion is 150,000 years (Stitzer *et al*., 2021). This landscape of old TE insertions are still highly polymorphic even among inbred breeding lines like the maize NAM parents, emphasized by the nearly 100 Mb variation in TE content we see in the NAM RILs. Here, we show that the cost of TEs in the maize genome is quantifiably small using large-scale fitness data sets from close to a million plants. While maize breeders have purged the most deleterious TE insertions, the cumulative load of small effects persists even in elite maize breeding populations as their selection coefficients are too small to effectively eradicate.

## Materials and Methods

Code to generate figures and analyses for this manuscript can be accessed at https://github.com/mcstitzer/maize_TE_load_fitness. The PHZ51 genome assembly is available upon request, and will be deposited at NCBI/ENA under Project PRJEB59044.

### Imputation of RIL haplotypes

To impute information from parental genome assemblies onto recombinant inbred lines (RILs), we use genotyping by sequencing (GBS) genotyping data of 6,624 replicates of 4,975 RILs derived from crosses between one of 25 inbred lines and the inbred line B73 (McMullen *et al*., 2009; Rodgers-Melnick *et al*., 2015, NCBI SRA SRP009896). We map reads from these individuals to a pangenome representation of parental assemblies using the Practical Haplotype Graph (PHG) (Bradbury *et al*., 2021), to capture parental haplotypes present in each RIL. Our reference pangenome database is Maize_1.0, with reference ranges based on B73v5 genes (Valdes Franco *et al*., 2020). We restrict haplotypes identified to those from any of B73 and 25 NAM parental assemblies from Hufford *et al*. (2021). For each RIL sample, we impute a path through the pangenome graph, allowing only haploid paths because these samples are inbred. This may ignore remaining residual heterozygosity, which is limited to less than 4% of markers in (McMullen *et al*., 2009). The individuals genotyped in this study have been self-fertilized for additional generations since McMullen *et al*. (2009), likely further increasing homozygosity. 1,649 individuals were genotyped in replicate – often biological replicates – when contamination or error was suspected or additional sequencing capacity was available. We select a single replicate to retain based on 1) if the two most likely parents imputed are those expected from the pedigree, 2) historical knowledge of GBS run failure, and if all else is equal, and 3) the replicate with highest read coverage.

Over several decades, mutation and contamination have generated true genetic differences between independent replicates of the B73 inbred line (Liang and Schnable, 2016). The B73 germplasm used to generate the NAM RILs comes from Major Goodman and is derived from the sub-line used by Pioneer Hi-Bred, while the stock used to assemble the genome is descended from USDA PI 550473 (Coe and Schaeffer, 2005), and further propagated by Michael McMullen. Regions of retained heterozygosity in the originally released B73 inbred line sorted into fixed homozygous differences between these lineages. Differences between genotyped regions of B73 have been noted before, including a region on chromosome 5 (Gore *et al*., 2009,, Supplemental Material). We take 38 genotyped replicates of B73 and 10 of each NAM parent (Romay *et al*., 2013), and impute their genotypes using the PHG as we did for each RIL. Regions with less than 60% (23 for B73, 6 for other parents) correctly assigned to the parental haplotype for the named genotyped sample are removed from future analysis. This removes a region on chromosome 5, found in none of the B73 individuals, as well as blocks on chromosomes 1 and 10. The PHG deals with large structural variation in a practical manner, collapsing regions such as inversion breakpoints into a single reference range, relative to the B73 allele, while positioning internal sequence at colinear regions. As there is segregating structural variation among the NAM parents, we removed 107 reference ranges with high variance in haplotype length among individuals (1 Mb difference between the maximum and minimum haplotype length).

For all figures, we show classification of NAM parents into categories of temperate, tropical, sweet, popcorn, and mixed germplasm from Flint-Garcia *et al*. (2005).

### TE Annotation

We project the TE annotation from each NAM parent onto haplotypes, summing contributions of TEs in total, each superfamily and family of TE, and knob, centromere, telomere, and ribosomal repeats. We used TE and repeat annotations updated from Hufford *et al*. (2021) and presented in Ou *et al*. (2022) (downloaded from https://de.cyverseorg/anon-files//iplant/home/shared/NAM/NAM_Canu1.8_TE_annotation_03032022/), collapsing by TE superfamily and repeat type. We summarized copies into superfamilies based on the Classification field in each parental gff, resulting in superfamilies DTA (Ac/Ds), DTC (CACTA), DTH (pIF/Harbinger), DTM (Mutator), DTT (Tc1/Mariner), DHH (Helitron), RIL (L1 LINE), RIT (RTE LINE), RIX (unknown LINE), RLC (Ty1/Copia), RLG (Ty3), RLX (Unknown LTR). We further assess contribution of TE families, focusing on the 170 families with greater than 10 Mb of sequence across all NAM parents. We combine LTR and internal regions of LTR retrotransposon records, and collapse different consensus copies of the same family into a single family identifier (e.g. ‘tekay_AC200856_6996’ and ‘tekay_AC211245_11065’ are both included in the Ty3 family ‘tekay’). For family-specific analyses, we remove families with names starting with ‘TE_’, due to inconsistencies between structural and homology-based superfamily assignment. In addition to TEs, we summarize the contribution of knob repeats, centromere repeats, telomere repeats, and ribosomal repeats to each haplotype. We repeat this process for each parental assembly, again, removing regions that cannot be genotyped or that differ between germplasm sources.

### TE characterization

We further assess features of TEs that have previously been tied to the deleterious impact of TEs on genes. We measure distance of each insertion to a core gene, as defined in Hufford *et al*. (2021). We sum TE base pairs within the gene, within 1 kilobase (kb) from the gene, from 1-5 kb from the gene, and greater than 5 kb. We assess the TE bp contributed by recent TE insertions, using a conservative metric that the TE copy has no divergence from the TE family consensus and is at least 500 bp, summarizing the youngest insertions in these maize genomes. We categorize all other TEs as ‘old’ insertions. As most TE families were originally defined based on the initial B73 genome assembly (Schnable *et al*., 2009), a majority of young TEs are inherited from the B73 parent. We identify TEs carrying an unmethylated region (UMRs) (Hufford *et al*., 2021), and calculate the amount of base pairs of TEs carrying UMRs in each RIL, and the amount of base pairs of TEs that lack an UMR.

### PHZ51 genome assembly and annotation

We assembled the genome of the former commercial tester line PHZ51, using PacBio CCS sequencing. We generated 212 Gb of sequence, and error-corrected these reads using mecat2 v20190314-8-gf54c542 (Xiao *et al*., 2017) with `CNS_OPTIONS=“-r 0.6 -a 1000 -c 4 -l 3000”`, selected the longest 40 reads when greater than 40x coverage with `CNS_OUTPUT_COVERAGE=40` and used a minimum read length of 2000. We then assembled these error-corrected reads using canu v 2.0 (Koren *et al*., 2017) with the `-trim-assemble` parameter, a kmer frequency threshold of `-ovlMerThreshold=500`, and `-genomeSize=2.5g`. This resulted in a 2 Gb assembly in 591 contigs, with an N50 of 3.5 Mb. To annotate TEs, we ran RepeatMasker v. 4.1.0, using `NAM.EDTA2.0.0.MTEC02052020.TElib.fa` as the repeat library, rmblastn as the search engine, and `-q -no_is` `-norna -nolow -div 40` parameters to match those used on the NAM maize assemblies. We summarized TEs from the gff3 output as above for other assemblies. For phenotypic analyses of hybrid maize, we add the genomic complement of TEs and repeats present in the PHZ51 parent to the RIL, creating a diploid genotypic value for the hybrid. To estimate the distance of TEs from genes, we projected the B73 gene models onto the PHZ51 assembly using Liftoff (Shumate and Salzberg, 2021).

### Phenotype Data

We used phenotypes collected from 1,723 hybrids of NAM RILs with a common PHZ51 tester parent from yield trials (Ramstein *et al*., 2020; Larsson *et al*., 2017). We use best linear unbiased estimator (BLUE) values from Ramstein *et al*. (2020) for days to silking (female flowering; N=1559), and a measure of grain yield incorporating female flowering as a fixed effect (N=1452), as yield is correlated to flowering time. The Hp301 popcorn family is not assayed in this experiment, so it is not present in results involving phenotypes.

## Associations

We associate genotypic descriptions of TE and repeat content with flowering time and grain yield phenotypes using a series of linear models. First, we associate the phenotype with genome size of each hybrid, in the form of *phenotype*_*i*_ ∼ *totalbp*_*i*_, where *i* indicates a NAM RIL hybrid. We next split this genome size phenotype into component parts - TE base pairs, knob, centromere, telomere, and ribosomal repeats, and nonTE-nonRepeat base pairs. We also include a fixed effect for the amount of base pairs of the B73 common parent to control for proportion ancestry of this common parent, resulting in a model of *phenotype*_*i*_ ∼ *TEbp*_*i*_ + *nonTEnonRepeatbp*_*i*_ + *knobbp*_*i*_ + *centromerebp*_*i*_ + *telomerebp*_*i*_ + *ribosomalbp*_*i*_ + *B*73*bp*_*i*_. As a complementary, more stringent correction for population structure, we incorporate three principal components (PCs) of a kinship matrix of the NAM RILs. We built this kinship matrix using SNPs from each chromosome. This model is thus *phenotype*_*i*_ ∼ *TEbp*_*i*_ + *nonTEnonRepeatbp*_*i*_ + *knobbp*_*i*_ + *centromerebp*_*i*_ + *telomerebp*_*i*_ + *ribosomalbp*_*i*_ + *PC*1_*i*_ + *PC*2_*i*_ + *PC*3_*i*_. To make use of the nested structure of the NAM population, we repeat the model using B73 bp control 24 times for each NAM family individually.

We test gene distance, TE age, and TE UMR presence in similar models. For gene distance, we fit a linear regression as *phenotype*_*i*_ ∼ *TEbpingene*_*i*_ + *TEbp*1*kbfromgene*_*i*_ + *TEbp*1*to*5*kbfromgene*_*i*_ + *TEbpgreaterthan*5*kbfromgene*_*i*_ + *B*73*bp*_*i*_ + *nonTEnonRepeatbp*_*i*_. For TE age, we fit a linear regression as *phenotype*_*i*_ ∼ *youngTEbp*_*i*_ + *oldT Ebp*_*i*_ + *B*73*bp*_*i*_ + *nonTEnonRepeatbp*_*i*_. For TE UMRs, we fit a linear regression as *phenotype*_*i*_ ∼ *TEbpwithUMR*_*i*_ + *TEbpwithoutUMR*_*i*_ + *B*73*bp*_*i*_ + *nonTEnonRepeatbp*_*i*_. To test the impact of individual TE families, we fit a linear regression in the form of *phenotype* ∼ *TEbpfam*1_*i*_ + *TEbpfam*2_*i*_ + … + *TEbpfam*170_*i*_ + *TEbpSmallerFams*_*i*_ + *B*73*bp* + *nonTEnonRepeatbp*_*i*_.

### Assumptions about yield

We aimed to convert our effect estimates from tonnes/hectare into easily interpretable values. These experiments were planted in two-row plots, with 40–80 plants per plot and 50,000–75,000 plants per hectare (Larsson *et al*., 2017; Ramstein *et al*., 2020). We thus use a mean value of 62,500 plants per hectare. An average ear of hybrid maize has 800 kernels, and each kernel weighs about 0.2 grams. By dividing our effect estimate using B73 bp as population structure correction by these values, we find 14.43 Mb of additional TE content decreases fitness by one kernel. An average TE fragment (across all genotypes) in Hufford *et al*. (2021) is 1599 base pairs. We consider the relative fitness between an individual with 800 kernels and 799 kernels, and divide the 14.4 Mb of TEs by their average length to count the 9005 TEs. This reduces to an average selection coefficient against a TE of 1.4e-7.

## ACKNOWLEDGEMENTS

M.C.S. was supported by NSF PRFB 1907343. M.C.S. thanks Joe Gage for reminding her how addition works, and Jeffrey Ross-Ibarra for supplying 14 Mb of TEs through the US Postal Service. We thank M. Cinta Romay and Peter Bradbury for advice and guidance about selecting genotypes of NAM RIL samples. We thank members of the Buckler and Feschotte Labs for comments and support.

