## Supplemental Information for "Transposable element abundance subtly contributes to lower fitness in maize"

### Supplementary Note 1: Supplemental Information

**Transposable element abundance subtly contributes to lower fitness in maize**

Stitzer, Khaipho-Burch, Hudson, Song, Valdez-Franco, Ramstein, Feschotte, Buckler (2023)

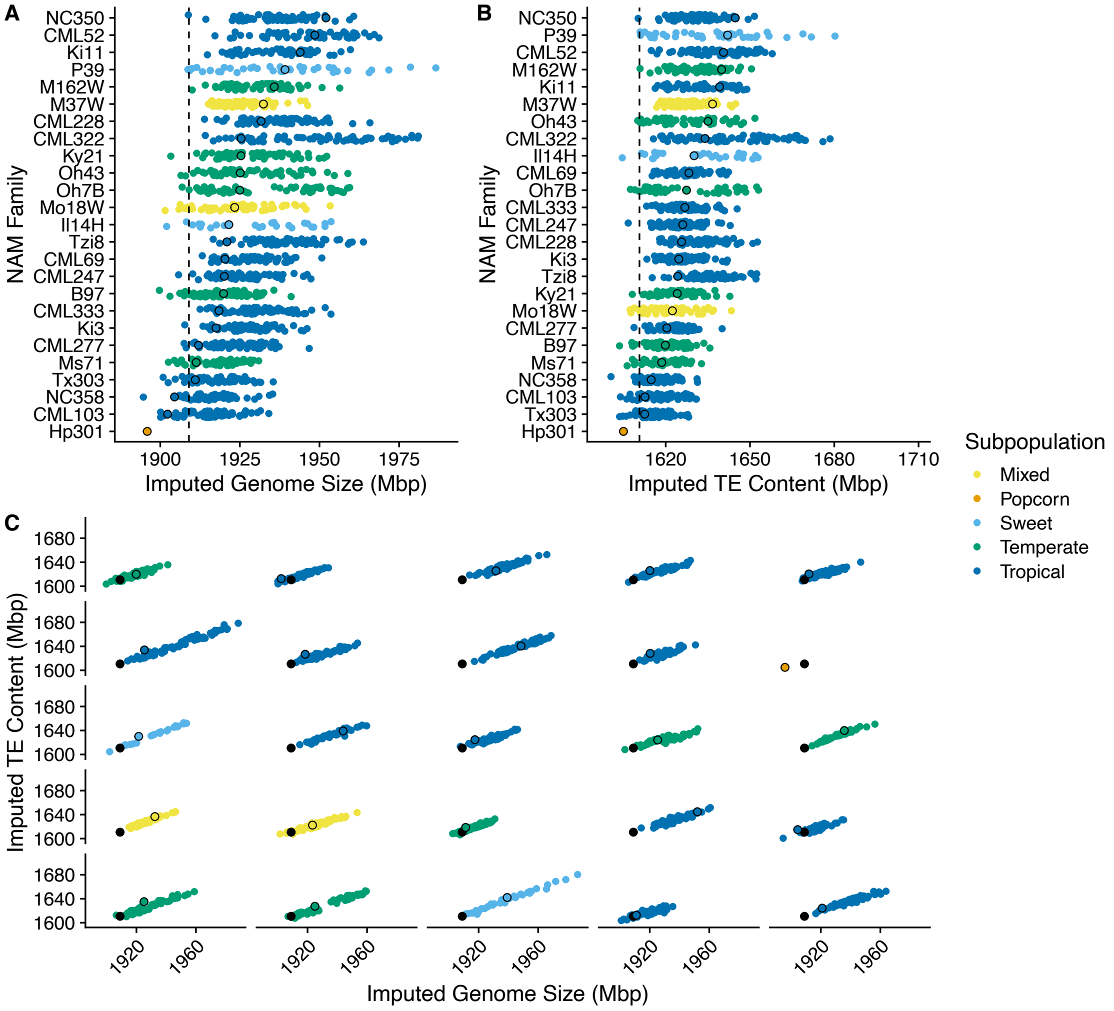


**Fig. S1. Imputed genome size and TE content for each NAM family, for individuals with phenotypes**. (A) Imputed genome size for each NAM family, (B) Imputed TE content for each NAM family, (C) Relationship between imputed genome content and imputed TE content in each NAM RIL. Each colored point reflects a NAM RIL, colored by maize subpopulation. Black outlined circle is the parental value calculated from the genome assembly, and black filled circles and dashed line are values for the B73 common parent. No Hp301 RILs were phenotyped, so only parental values are plotted.

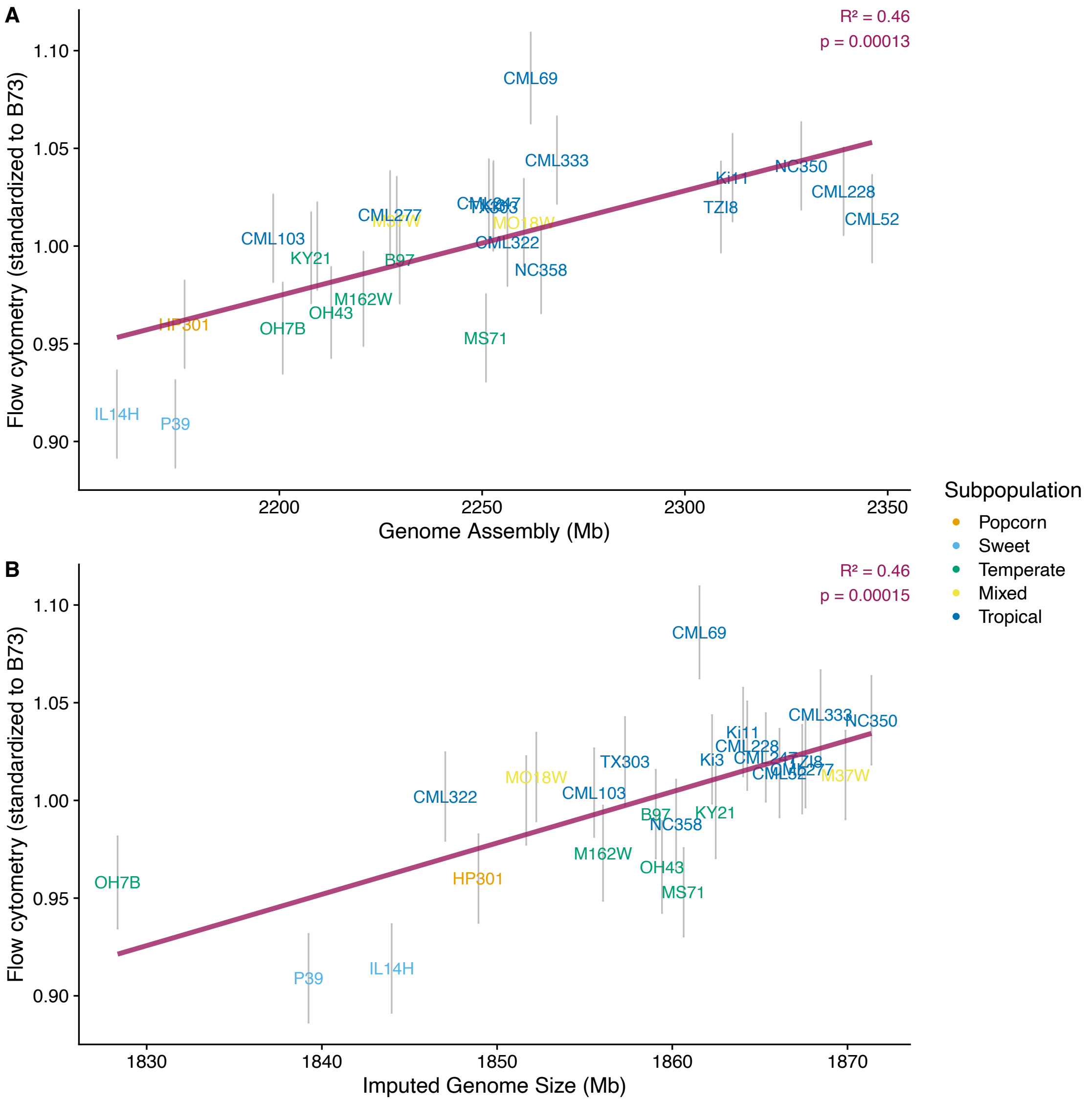


**Fig. S2. Association of genome assembly size and imputed genome size with flow cytometry estimates of genome size**. (A) Size of genome assembly vs flow cytometry values standardized to B73, (B) Imputed genome size vs flow cytometry values standardized to B73. Standardized flow cytometry values from Chia *et al.* (2012) Each text label shows a NAM parent, colored by maize subpopulation, with standard errors shown in gray line. Purple lines show linear regression.

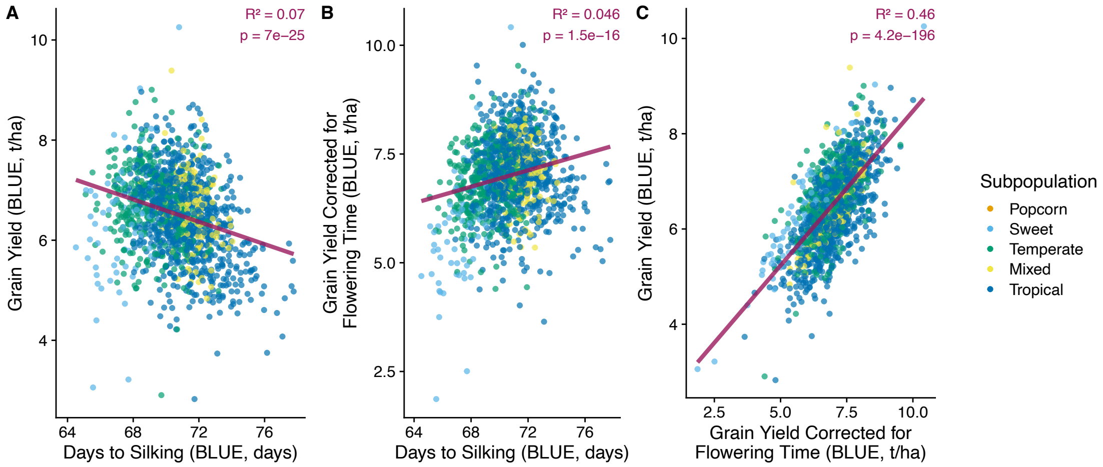


**Fig. S3. Associations between phenotypes**. (A) Grain Yield vs Days to Silking, (B) Grain Yield adjusted for DTS vs Days to Silking, (C) Raw Grain Yield vs Grain Yield corrected for DTS. Each colored point reflects a NAM RIL, colored by maize subpopulation. Lines show linear regression.

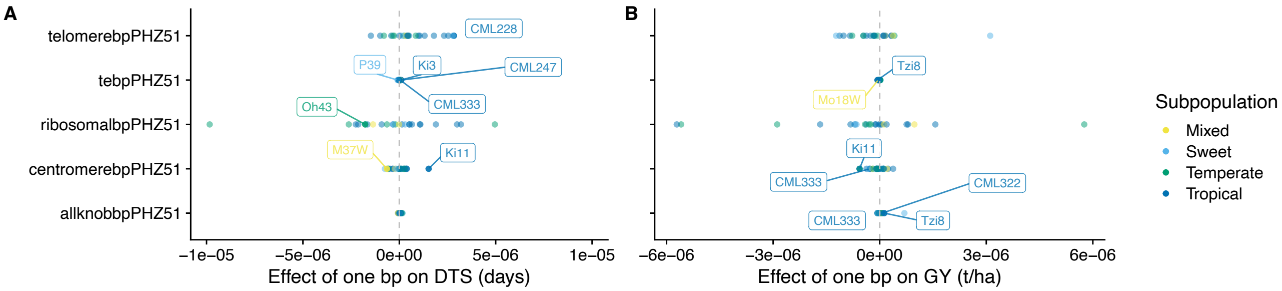


**Fig. S4. Phenotype associations with genomic repeats, for each NAM family**. (A) Female flowering time (Days to Silking), (B) Grain yield. Y axis shows different genomic repeat category, x axis shows estimated effect. Each colored point reflects the regression coefficient of a NAM family, colored by maize subpopulation. Families that are significantly associated are plotted in a darker color, and labeled with their family name. Model summaries in Supp. Tables S3 and S4.

| | DTS | GY |
|---|---|---|
| (Intercept) | $7.31 \times 10^{1}$*** | $4.17 \times 10^{1}$*** |
| TE bp | $-3.19 \times 10^{-8}$*** | $-1.29 \times 10^{-8}$*** |
| nonTE, nonRepeat bp | $2.11 \times 10^{-7}$*** | $2.05 \times 10^{-8}$+ |
| Knob bp | $1.04 \times 10^{-7}$*** | $1.25 \times 10^{-8}$+ |
| Centromere bp | $2.01 \times 10^{-7}$* | $-2.38 \times 10^{-8}$ |
| Telomere bp | $-6.69 \times 10^{-8}$ | $7.82 \times 10^{-8}$ |
| Ribosomal bp | $-4.66 \times 10^{-9}$ | $3.69 \times 10^{-7}$* |
| PC1 | $6.38 \times 10^{1}$*** | $1.14 \times 10^{1}$*** |
| PC2 | 1.51 | $-7.15 \times 10^{-1}$ |
| PC3 | 9.13** | 7.38*** |
| Num.Obs. | 1558 | 1451 |
| R2 | 0.333 | 0.068 |
| R2 Adj. | 0.329 | 0.062 |

+ $p < 0.1$, * $p < 0.05$, ** $p < 0.01$, *** $p < 0.001$

**Table S1.** Effect sizes of the relationship between TE content, genomic repeats, principal components (PCs) of a kinship matrix with flowering time (DTS) and grain yield (GY).

| | DTS | GY |
|---|---|---|
| (Intercept) | $6.01 \times 10^{1}$*** | 4.48 |
| DHH (Helitron) bp | $-5.36 \times 10^{-7}$*** | $-2.81 \times 10^{-7}$*** |
| DTA (Ac/Ds) bp | $-8.40 \times 10^{-7}$* | $5.36 \times 10^{-8}$ |
| DTC (En/Spm/CACTA) bp | $5.03 \times 10^{-7}$*** | $-4.08 \times 10^{-8}$ |
| DTH (pIF/Harbinger) bp | $-8.97 \times 10^{-7}$+ | $6.42 \times 10^{-7}$* |
| DTM (Mutator) bp | $-4.68 \times 10^{-7}$*** | $-6.97 \times 10^{-8}$ |
| DTT (Tc1/Mariner) bp | $-1.53 \times 10^{-6}$ | $-1.29 \times 10^{-6}$ |
| RIL (L1) bp | $-1.76 \times 10^{-6}$*** | $-7.65 \times 10^{-7}$** |
| RIT (RTE) bp | $-5.06 \times 10^{-6}$** | $-7.90 \times 10^{-7}$ |
| RLC (Ty1/Copia) bp | $9.92 \times 10^{-8}$*** | $6.30 \times 10^{-8}$*** |
| RLG (Ty3) bp | $-3.14 \times 10^{-8}$* | $-2.95 \times 10^{-8}$*** |
| RLX (Unclassified LTR) bp | $-7.31 \times 10^{-8}$ | $5.76 \times 10^{-8}$ |
| nonTE, nonRepeat bp | $2.09 \times 10^{-7}$*** | $6.83 \times 10^{-9}$ |
| Knob bp | $8.46 \times 10^{-8}$*** | $1.13 \times 10^{-8}$ |
| Centromere bp | $4.39 \times 10^{-8}$ | $-2.79 \times 10^{-8}$ |
| Telomere bp | $2.70 \times 10^{-7}$ | $8.38 \times 10^{-8}$ |
| Ribosomal bp | $1.61 \times 10^{-7}$ | $2.36 \times 10^{-7}$ |
| B73 bp | $-2.48 \times 10^{-11}$ | $6.92 \times 10^{-11}$ |
| PC1 | $4.55 \times 10^{1}$*** | 2.63 |
| PC2 | 6.39+ | $-1.63 \times 10^{-1}$ |
| PC3 | 5.94+ | 4.63** |
| Num.Obs. | 1558 | 1451 |
| R2 | 0.373 | 0.109 |
| R2 Adj. | 0.364 | 0.097 |
| AIC | 5930.0 | 3571.7 |
| BIC | 6047.7 | 3687.9 |
| Log.Lik. | −2942.984 | −1763.867 |
| RMSE | 1.60 | 0.82 |

+ $p < 0.1$, * $p < 0.05$, ** $p < 0.01$, *** $p < 0.001$

**Table S2.** Effect sizes of the relationship between TE content by superfamily, genomic repeats and flowering time (DTS) and grain yield (GY).

| | DTS_B97 | DTS_CML103 | DTS_CML228 | DTS_CML247 | DTS_CML277 | DTS_CML322 | DTS_CML333 | DTS_CML52 |
|---|---|---|---|---|---|---|---|---|
| (Intercept) | $-1.16\times10^{2}$ | $-9.40\times10^{1}$ | $-1.06\times10^{2}$ | $-3.60\times10^{2}$** | $5.90\times10^{1}$ | $3.21\times10^{1}$ | $-2.25\times10^{2}$* | $-1.42\times10^{2}$+ |
| TE bp | $2.49\times10^{-10}$ | $2.52\times10^{-8}$ | $1.15\times10^{-8}$ | $1.02\times10^{-7}$** | $6.52\times10^{-9}$ | $-1.39\times10^{-9}$ | $4.94\times10^{-8}$* | $5.68\times10^{-8}$+ |
| Knob bp | $-1.15\times10^{-8}$ | $1.63\times10^{-8}$ | $1.16\times10^{-8}$ | $-4.60\times10^{-8}$ | $9.70\times10^{-8}$ | $4.62\times10^{-8}$ | $-6.19\times10^{-8}$ | $1.09\times10^{-7}$+ |
| Centromere bp | $-2.77\times10^{-8}$ | $3.77\times10^{-7}$ | $3.85\times10^{-7}$ | $-4.91\times10^{-7}$+ | $-3.25\times10^{-7}$ | $-3.58\times10^{-7}$ | $2.88\times10^{-7}$ | $-5.73\times10^{-7}$ |
| Telomere bp | $-2.66\times10^{-7}$ | $1.06\times10^{-6}$ | $2.83\times10^{-6}$* | $9.92\times10^{-7}$ | $1.31\times10^{-6}$ | $-4.16\times10^{-7}$ | $-9.90\times10^{-7}$ | $1.80\times10^{-6}$ |
| Ribosomal bp | $4.96\times10^{-6}$ | $5.25\times10^{-7}$ | $3.00\times10^{-6}$ | $-3.25\times10^{-7}$ | $-9.15\times10^{-7}$ | $1.43\times10^{-7}$ | $1.09\times10^{-6}$ | $1.90\times10^{-6}$ |
| nonTE, nonRepeat bp | $3.44\times10^{-7}$** | $1.31\times10^{-7}$ | $2.35\times10^{-7}$ | $1.45\times10^{-7}$ | $-1.97\times10^{-8}$ | $9.26\times10^{-8}$ | $2.37\times10^{-7}$+ | $1.10\times10^{-8}$ |
| Num.Obs. | 66 | 67 | 77 | 72 | 65 | 73 | 66 | 61 |
| R2 | 0.143 | 0.111 | 0.155 | 0.196 | 0.053 | 0.066 | 0.264 | 0.250 |
| R2 Adj. | 0.056 | 0.022 | 0.082 | 0.121 | −0.045 | −0.019 | 0.189 | 0.167 |

| | DTS_CML69 | DTS_Il14H | DTS_Ki11 | DTS_Ki3 | DTS_Ky21 | DTS_M162W | DTS_M37W | DTS_Mo18W |
|---|---|---|---|---|---|---|---|---|
| (Intercept) | $-4.24\times10^{1}$ | $-4.89\times10^{1}$ | $4.96\times10^{-1}$ | $-4.24\times10^{1}$ | $1.42\times10^{2}$ | $1.42\times10^{2}$+ | $1.37\times10^{2}$+ | $-3.67\times10^{1}$ |
| TE bp | $-5.43\times10^{-9}$ | $4.62\times10^{-8}$+ | $-3.74\times10^{-8}$ | $5.56\times10^{-8}$* | $-2.39\times10^{-8}$ | $1.85\times10^{-8}$ | $2.37\times10^{-8}$ | $3.51\times10^{-8}$ |
| Knob bp | $2.60\times10^{-8}$ | $1.68\times10^{-7}$ | $6.52\times10^{-8}$ | $4.01\times10^{-8}$ | $1.47\times10^{-7}$ | $6.99\times10^{-8}$ | $-7.80\times10^{-8}$ | $-2.83\times10^{-8}$ |
| Centromere bp | $2.43\times10^{-7}$ | $-7.50\times10^{-7}$+ | $1.52\times10^{-6}$*** | $1.38\times10^{-8}$ | $-5.81\times10^{-7}$ | $2.14\times10^{-7}$ | $-6.44\times10^{-7}$* | $-2.27\times10^{-7}$ |
| Telomere bp | $-1.48\times10^{-6}$ | $2.77\times10^{-6}$+ | $2.54\times10^{-6}$ | $5.03\times10^{-7}$ | $2.47\times10^{-7}$ | $-3.44\times10^{-7}$ | $4.00\times10^{-7}$ | $3.43\times10^{-7}$ |
| Ribosomal bp | $-2.13\times10^{-6}$ | $3.05\times10^{-7}$ | $-1.66\times10^{-6}$ | $1.07\times10^{-6}$ | $-9.84\times10^{-6}$+ | $-2.05\times10^{-7}$ | $-1.10\times10^{-8}$ | $-1.37\times10^{-6}$ |
| nonTE, nonRepeat bp | $2.76\times10^{-7}$+ | $-1.02\times10^{-7}$ | $3.69\times10^{-7}$ | $-1.82\times10^{-7}$ | $8.70\times10^{-8}$* | $-2.84\times10^{-7}$ | $-2.83\times10^{-7}$+ | $-2.12\times10^{-8}$ |
| Num.Obs. | 67 | 46 | 53 | 74 | 70 | 67 | 68 | 62 |
| R2 | 0.090 | 0.155 | 0.318 | 0.107 | 0.135 | 0.059 | 0.100 | 0.099 |
| R2 Adj. | −0.001 | 0.026 | 0.229 | 0.027 | 0.052 | −0.035 | 0.012 | 0.000 |

| | DTS_Ms71 | DTS_NC350 | DTS_NC358 | DTS_Oh43 | DTS_Oh7B | DTS_P39 | DTS_Tx303 | DTS_Tzi8 |
|---|---|---|---|---|---|---|---|---|
| (Intercept) | $1.13\times10^{2}$ | $-1.23\times10^{2}$ | $-6.42\times10^{1}$ | $1.11\times10^{2}$* | $1.08\times10^{2}$* | $1.87\times10^{2}$*** | $1.60\times10^{2}$ | $-5.00\times10^{1}$ |
| TE bp | $-1.28\times10^{-8}$ | $2.40\times10^{-8}$ | $2.04\times10^{-9}$ | $-2.85\times10^{-8}$ | $-2.07\times10^{-8}$ | $-8.42\times10^{-8}$** | $-2.63\times10^{-8}$ | $-8.27\times10^{-10}$ |
| Knob bp | $2.47\times10^{-8}$ | $8.64\times10^{-8}$ | $4.88\times10^{-8}$ | $2.98\times10^{-8}$ | $2.65\times10^{-8}$ | $1.15\times10^{-8}$ | $3.41\times10^{-8}$ | $-8.10\times10^{-9}$ |
| Centromere bp | $1.28\times10^{-8}$ | $-3.92\times10^{-8}$ | $4.70\times10^{-8}$ | $2.99\times10^{-7}$ | $9.56\times10^{-8}$ | $-1.67\times10^{-7}$ | $1.22\times10^{-7}$ | $3.78\times10^{-7}$+ |
| Telomere bp | $4.55\times10^{-7}$ | $2.32\times10^{-6}$ | $4.02\times10^{-7}$ | $-7.99\times10^{-7}$ | $8.51\times10^{-7}$ | $1.07\times10^{-7}$ | $7.86\times10^{-9}$ | $4.46\times10^{-7}$ |
| Ribosomal bp | $-6.40\times10^{-7}$ | $5.20\times10^{-7}$ | $3.20\times10^{-6}$ | $-1.77\times10^{-6}$* | $-2.62\times10^{-6}$ | $-3.31\times10^{-7}$ | $6.58\times10^{-7}$ | $-2.26\times10^{-6}$ |
| nonTE, nonRepeat bp | $3.08\times10^{-10}$ | $2.00\times10^{-7}$ | $2.29\times10^{-7}$+ | $1.20\times10^{-7}$ | $7.88\times10^{-8}$ | $3.67\times10^{-7}$+ | $8.25\times10^{-10}$ | $2.47\times10^{-7}$+ |
| Num.Obs. | 58 | 67 | 65 | 63 | 64 | 58 | 65 | 67 |
| R2 | 0.017 | 0.128 | 0.103 | 0.172 | 0.089 | 0.300 | 0.026 | 0.235 |
| R2 Adj. | −0.098 | 0.041 | 0.010 | 0.083 | −0.007 | 0.218 | −0.075 | 0.158 |

+ $p < 0.1$, * $p < 0.05$, ** $p < 0.01$, *** $p < 0.001$

**Table S3.** Effect sizes of the relationship between TE content by superfamily, genomic repeats and flowering time (DTS), for each NAM family.

| | GY_B97 | GY_CML103 | GY_CML228 | GY_CML247 | GY_CML277 | GY_CML322 | GY_CML333 | GY_CML52 |
|---|---|---|---|---|---|---|---|---|
| (Intercept) | $-1.18 \times 10^{1}$ | $-5.53 \times 10^{1}$ | $9.35 \times 10^{1}$+ | $2.18 \times 10^{2}$** | $4.91 \times 10^{1}$ | $4.89 \times 10^{1}$* | $-1.94 \times 10^{1}$ | $4.43 \times 10^{1}$ |
| tebpPHZ51 | $1.01 \times 10^{-8}$ | $-2.36 \times 10^{-8}$ | $-1.40 \times 10^{-8}$ | $-3.90 \times 10^{-8}$+ | $1.02 \times 10^{-8}$ | $-7.79 \times 10^{-9}$ | $1.32 \times 10^{-8}$ | $-1.44 \times 10^{-8}$ |
| allknobbpPHZ51 | $2.00 \times 10^{-8}$ | $-1.91 \times 10^{-8}$ | $2.05 \times 10^{-8}$ | $3.35 \times 10^{-9}$ | $-5.46 \times 10^{-8}$+ | $6.71 \times 10^{-8}$** | $-6.43 \times 10^{-8}$* | $1.63 \times 10^{-9}$ |
| centromerebpPHZ51 | $-5.17 \times 10^{-8}$ | $-3.71 \times 10^{-7}$+ | $-7.87 \times 10^{-8}$ | $2.40 \times 10^{-7}$ | $-1.69 \times 10^{-9}$ | $-4.11 \times 10^{-8}$ | $-2.76 \times 10^{-7}$* | $-2.27 \times 10^{-8}$ |
| telomerebpPHZ51 | $-1.77 \times 10^{-7}$ | $3.26 \times 10^{-7}$ | $-3.73 \times 10^{-7}$ | $-4.55 \times 10^{-7}$ | $-2.59 \times 10^{-7}$ | $1.07 \times 10^{-7}$ | $1.24 \times 10^{-7}$ | $-8.37 \times 10^{-7}$ |
| ribosomalbpPHZ51 | $-2.89 \times 10^{-6}$ | $7.97 \times 10^{-7}$ | $-1.07 \times 10^{-7}$ | $1.68 \times 10^{-8}$ | $-3.77 \times 10^{-7}$ | $-6.86 \times 10^{-7}$ | $-2.43 \times 10^{-8}$ | $5.18 \times 10^{-8}$ |
| nontenonrepeatbpPHZ51 | $-1.84 \times 10^{-8}$ | $2.97 \times 10^{-7}$* | $-6.97 \times 10^{-8}$ | $-1.47 \times 10^{-7}$ | $-1.50 \times 10^{-7}$ | $-2.76 \times 10^{-8}$ | $-3.04 \times 10^{-8}$ | $3.14 \times 10^{-8}$ |
| Num.Obs. | 66 | 67 | 72 | 70 | 63 | 72 | 65 | 59 |
| R2 | 0.019 | 0.179 | 0.060 | 0.146 | 0.111 | 0.179 | 0.177 | 0.039 |
| R2 Adj. | −0.081 | 0.097 | −0.027 | 0.065 | 0.016 | 0.104 | 0.092 | −0.072 |

| | GY_CML69 | GY_Il14H | GY_Ki11 | GY_Ki3 | GY_Ky21 | GY_M162W | GY_M37W | GY_Mo18W |
|---|---|---|---|---|---|---|---|---|
| (Intercept) | $3.39 \times 10^{1}$ | $-1.77 \times 10^{2}$ | $3.90 \times 10^{1}$ | $7.27 \times 10^{1}$ | $-4.64 \times 10^{1}$ | $1.55 \times 10^{1}$ | $1.17 \times 10^{2}$* | $1.79 \times 10^{2}$*** |
| tebpPHZ51 | $-6.83 \times 10^{-10}$ | $2.28 \times 10^{-8}$ | $-1.11 \times 10^{-8}$ | $-2.09 \times 10^{-9}$ | $1.45 \times 10^{-8}$ | $2.30 \times 10^{-9}$ | $-2.32 \times 10^{-8}$ | $-4.41 \times 10^{-8}$* |
| allknobbpPHZ51 | $1.74 \times 10^{-8}$ | $6.96 \times 10^{-7}$ | $-3.28 \times 10^{-8}$ | $-2.03 \times 10^{-8}$ | $1.84 \times 10^{-8}$ | $7.65 \times 10^{-8}$ | $2.47 \times 10^{-8}$ | $5.48 \times 10^{-8}$ |
| centromerebpPHZ51 | $9.50 \times 10^{-8}$ | $1.03 \times 10^{-7}$ | $-5.67 \times 10^{-7}$*** | $5.24 \times 10^{-8}$ | $-5.63 \times 10^{-7}$ | $-9.11 \times 10^{-8}$ | $-2.09 \times 10^{-7}$ | $2.05 \times 10^{-7}$ |
| telomerebpPHZ51 | $-1.02 \times 10^{-6}$ | $3.11 \times 10^{-6}$ | $-1.33 \times 10^{-7}$ | $3.33 \times 10^{-7}$ | $-4.61 \times 10^{-7}$ | $4.15 \times 10^{-7}$ | $1.55 \times 10^{-7}$ | $3.71 \times 10^{-7}$ |
| ribosomalbpPHZ51 | $7.59 \times 10^{-7}$ | $-7.14 \times 10^{-7}$ | $-1.27 \times 10^{-7}$ | $-1.68 \times 10^{-6}$+ | $-5.59 \times 10^{-6}$ | $-2.48 \times 10^{-7}$ | $9.77 \times 10^{-7}$ | $1.17 \times 10^{-7}$ |
| nontenonrepeatbpPHZ51 | $-5.07 \times 10^{-8}$ | $1.49 \times 10^{-7}$ | $3.24 \times 10^{-8}$ | $-1.08 \times 10^{-7}$ | $4.83 \times 10^{-8}$ | $-3.64 \times 10^{-8}$ | $-5.71 \times 10^{-8}$ | $-3.95 \times 10^{-8}$ |
| Num.Obs. | 64 | 24 | 52 | 69 | 56 | 67 | 67 | 61 |
| R2 | 0.082 | 0.228 | 0.323 | 0.094 | 0.120 | 0.052 | 0.113 | 0.182 |
| R2 Adj. | −0.014 | −0.045 | 0.232 | 0.006 | 0.013 | −0.043 | 0.025 | 0.091 |

| | GY_Ms71 | GY_NC350 | GY_NC358 | GY_Oh43 | GY_Oh7B | GY_P39 | GY_Tx303 | GY_Tzi8 |
|---|---|---|---|---|---|---|---|---|
| (Intercept) | $3.87 \times 10^{1}$ | $4.88 \times 10^{1}$ | $7.14 \times 10^{1}$ | $-9.48$ | $-2.26 \times 10^{1}$ | $4.01 \times 10^{1}$ | $5.08 \times 10^{1}$ | $1.03 \times 10^{2}$ |
| tebpPHZ51 | $1.91 \times 10^{-8}$ | $-1.63 \times 10^{-8}$ | $-9.09 \times 10^{-9}$ | $-4.55 \times 10^{-9}$ | $-1.42 \times 10^{-8}$ | $-5.01 \times 10^{-8}$+ | $-1.09 \times 10^{-8}$ | $-6.25 \times 10^{-8}$* |
| allknobbpPHZ51 | $-4.94 \times 10^{-8}$ | $-1.80 \times 10^{-8}$ | $5.49 \times 10^{-8}$ | $2.85 \times 10^{-8}$ | $-6.12 \times 10^{-9}$ | $-5.58 \times 10^{-8}$ | $-2.22 \times 10^{-8}$ | $1.30 \times 10^{-7}$** |
| centromerebpPHZ51 | $-1.86 \times 10^{-7}$ | $-9.98 \times 10^{-8}$ | $-8.48 \times 10^{-8}$ | $-6.83 \times 10^{-8}$ | $1.04 \times 10^{-7}$ | $-3.18 \times 10^{-7}$ | $1.21 \times 10^{-7}$ | $3.78 \times 10^{-7}$+ |
| telomerebpPHZ51 | $-1.54 \times 10^{-7}$ | $-1.13 \times 10^{-6}$ | $1.61 \times 10^{-7}$ | $-7.73 \times 10^{-7}$ | $-6.19 \times 10^{-8}$ | $-1.23 \times 10^{-6}$ | $-1.64 \times 10^{-7}$ | $9.60 \times 10^{-8}$ |
| ribosomalbpPHZ51 | $-3.00 \times 10^{-7}$ | $-5.71 \times 10^{-6}$ | $1.57 \times 10^{-6}$ | $-4.14 \times 10^{-7}$ | $5.76 \times 10^{-6}$+ | $-6.51 \times 10^{-7}$ | $1.95 \times 10^{-7}$ | $-8.21 \times 10^{-7}$ |
| nontenonrepeatbpPHZ51 | $-1.91 \times 10^{-7}$+ | $7.34 \times 10^{-8}$ | $-7.51 \times 10^{-8}$ | $7.00 \times 10^{-8}$ | $1.26 \times 10^{-7}$ | $3.08 \times 10^{-7}$ | $-1.23 \times 10^{-8}$ | $2.40 \times 10^{-7}$+ |
| Num.Obs. | 55 | 64 | 62 | 58 | 57 | 39 | 64 | 61 |
| R2 | 0.104 | 0.108 | 0.092 | 0.098 | 0.106 | 0.181 | 0.021 | 0.216 |
| R2 Adj. | −0.008 | 0.014 | −0.008 | −0.009 | −0.001 | 0.027 | −0.082 | 0.128 |

+ $p < 0.1$, * $p < 0.05$, ** $p < 0.01$, *** $p < 0.001$

**Table S4.** Effect sizes of the relationship between TE content by superfamily, genomic repeats and grain yield (GY), for each NAM family.

| | GY | DTS |
|---|---|---|
| (Intercept) | $-5.15 \times 10^{1}$** | $-1.57 \times 10^{2}$*** |
| Within gene TE base pairs | $3.44 \times 10^{-7}$*** | $7.50 \times 10^{-7}$*** |
| One kb from gene TE base pairs | $-1.48 \times 10^{-7}$* | $-1.72 \times 10^{-6}$*** |
| One to five kb from gene TE base pairs | $1.12 \times 10^{-7}$*** | $1.74 \times 10^{-7}$** |
| Greater than five kb from gene TE base pairs | $3.37 \times 10^{-9}$ | $9.32 \times 10^{-8}$*** |
| B73 bp | $2.45 \times 10^{-10}$* | $-3.89 \times 10^{-10}$+ |
| Num.Obs. | 1454 | 1561 |
| R2 | 0.040 | 0.235 |
| R2 Adj. | 0.037 | 0.233 |

+ $p < 0.1$, * $p < 0.05$, ** $p < 0.01$, *** $p < 0.001$

**Table S5.** Effect sizes of the relationship between TE content at different gene distances and flowering time (DTS) or grain yield (GY).

| | GY | DTS |
|---|---|---|
| (Intercept) | $-9.03$ | $-2.33 \times 10^{2}$*** |
| Recent TE base pairs | $-1.78 \times 10^{-6}$*** | $-5.30 \times 10^{-6}$*** |
| Older TE base pairs | $5.10 \times 10^{-9}$ | $8.72 \times 10^{-8}$*** |
| B73 bp | $1.03 \times 10^{-9}$*** | $1.93 \times 10^{-9}$*** |
| Num.Obs. | 1454 | 1561 |
| R2 | 0.016 | 0.138 |
| R2 Adj. | 0.014 | 0.136 |

+ $p < 0.1$, * $p < 0.05$, ** $p < 0.01$, *** $p < 0.001$

**Table S6.** Effect sizes of the relationship between recent (<30,000 years) and older (>30,000 years) TE base pairs and flowering time (DTS) or grain yield (GY).

| | GY | DTS |
|---|---|---|
| (Intercept) | $-6.45$ | $-7.59 \times 10^{1}$*** |
| TE without UMR base pairs | $8.74 \times 10^{-9}$* | $9.24 \times 10^{-8}$*** |
| TE with UMR base pairs | $1.09 \times 10^{-9}$ | $1.20 \times 10^{-7}$*** |
| teha$b73bp | $1.56 \times 10^{-11}$ | $-1.21 \times 10^{-9}$*** |
| Num.Obs. | 1454 | 1561 |
| R2 | 0.005 | 0.121 |
| R2 Adj. | 0.003 | 0.119 |

+ $p < 0.1$, * $p < 0.05$, ** $p < 0.01$, *** $p < 0.001$

**Table S7.** Effect sizes of the relationship between base pairs of TEs with unmethylated regions (UMRs) and TEs without UMRs and flowering time (DTS) or grain yield (GY).

| | GY | DTS |
|---|---|---|
| (Intercept) | $4.27 \times 10^{1}$*** | $9.65 \times 10^{1}$*** |
| b73bp | $-2.16 \times 10^{-10}$ | $-1.49 \times 10^{-9}$*** |
| smallerFamTEbp | $-9.05 \times 10^{-9}$*** | $-7.96 \times 10^{-9}$+ |
| huck | $-4.32 \times 10^{-8}$* | $-3.72 \times 10^{-8}$ |
| cinful | $3.28 \times 10^{-8}$ | $6.61 \times 10^{-8}$ |
| opie | $1.52 \times 10^{-8}$ | $-4.22 \times 10^{-8}$ |
| ji | $5.37 \times 10^{-8}$ | $1.05 \times 10^{-7}$ |
| flip | $-6.48 \times 10^{-8}$+ | $-2.30 \times 10^{-8}$ |
| xilon | $2.65 \times 10^{-9}$ | $2.48 \times 10^{-7}$** |
| chr6_P_87978263 | $1.33 \times 10^{-7}$** | $-5.79 \times 10^{-8}$ |
| gyma | $-3.93 \times 10^{-8}$ | $-5.83 \times 10^{-8}$ |
| prem1 | $4.02 \times 10^{-8}$ | $-2.46 \times 10^{-7}$** |
| grande | $1.37 \times 10^{-8}$ | $1.42 \times 10^{-7}$ |
| doke | $1.26 \times 10^{-7}$* | $-1.25 \times 10^{-8}$ |
| giepum | $3.09 \times 10^{-8}$ | $1.20 \times 10^{-7}$ |
| milt | $-3.10 \times 10^{-7}$* | $2.76 \times 10^{-8}$ |
| iteki | $-1.67 \times 10^{-7}$ | $5.00 \times 10^{-8}$ |
| mada | $2.93 \times 10^{-7}$* | $3.87 \times 10^{-7}$+ |
| uwum | $1.00 \times 10^{-8}$ | $1.29 \times 10^{-7}$ |
| ruda | $-9.45 \times 10^{-8}$ | $3.19 \times 10^{-7}$ |
| dagaf | $8.02 \times 10^{-8}$ | $4.33 \times 10^{-7}$+ |
| chr10_P_98050947 | $-3.60 \times 10^{-7}$* | $-6.85 \times 10^{-7}$* |
| tekay | $-5.97 \times 10^{-8}$ | $1.42 \times 10^{-7}$ |
| chr4_P_242404877 | $-1.05 \times 10^{-7}$ | $-7.71 \times 10^{-7}$* |
| chr9_P_79032327 | $-7.13 \times 10^{-9}$ | $-3.29 \times 10^{-7}$ |
| chr6_D_43881252 | $5.84 \times 10^{-9}$ | $-4.71 \times 10^{-7}$ |
| DTC_ZM00081_consensus | $-4.57 \times 10^{-7}$* | $-4.41 \times 10^{-7}$ |
| chr2_D_224980689 | $3.22 \times 10^{-7}$ | $3.84 \times 10^{-7}$ |
| chr9_D_120499978 | $-1.08 \times 10^{-7}$ | $2.96 \times 10^{-7}$ |
| nida | $-3.29 \times 10^{-7}$+ | $-1.29 \times 10^{-7}$ |
| leviathan | $2.67 \times 10^{-7}$ | $7.51 \times 10^{-7}$ |
| chr9_D_32444039 | $-3.23 \times 10^{-7}$ | $-6.67 \times 10^{-7}$ |
| ansuya | $1.81 \times 10^{-7}$ | $-4.91 \times 10^{-7}$+ |
| chr5_D_119412279 | $5.51 \times 10^{-8}$ | $3.94 \times 10^{-7}$ |
| DTC_ZM00018_consensus | $-2.82 \times 10^{-7}$ | $3.51 \times 10^{-7}$ |
| chr4_P_103002213 | $-4.08 \times 10^{-7}$ | $6.77 \times 10^{-7}$ |
| wiwa | $-3.04 \times 10^{-7}$ | $1.20 \times 10^{-7}$ |
| chr3_P_39978843 | $-5.69 \times 10^{-7}$ | $-3.25 \times 10^{-6}$*** |
| chr10_D_107123494 | $-3.36 \times 10^{-7}$ | $6.06 \times 10^{-7}$ |
| chr4_D_40959532 | $-7.14 \times 10^{-8}$ | $-7.34 \times 10^{-7}$+ |
| chr1_D_275633391 | $1.62 \times 10^{-7}$ | $4.42 \times 10^{-7}$ |
| CRM2 | $-1.19 \times 10^{-7}$ | $-2.42 \times 10^{-7}$+ |
| nihep | $1.31 \times 10^{-7}$ | $4.10 \times 10^{-7}$ |
| gori | $-3.48 \times 10^{-7}$ | $1.22 \times 10^{-6}$* |
| chr6_D_62813001 | $1.41 \times 10^{-7}$ | $1.81 \times 10^{-6}$*** |
| chr5_P_76002780 | $-4.24 \times 10^{-7}$ | $5.76 \times 10^{-7}$ |
| DTC_ZM00004_consensus | $-6.37 \times 10^{-7}$* | $-2.36 \times 10^{-6}$*** |
| gudyeg | $2.06 \times 10^{-8}$ | $7.68 \times 10^{-7}$ |
| gunu | $2.07 \times 10^{-7}$ | $-6.89 \times 10^{-7}$ |
| naadira | $-5.64 \times 10^{-7}$ | $-1.42 \times 10^{-6}$* |
| ubow | $-2.58 \times 10^{-7}$ | $-3.02 \times 10^{-6}$** |
| chr7_D_125343800 | $3.50 \times 10^{-7}$ | $-5.48 \times 10^{-7}$ |
| odoj | $-7.17 \times 10^{-7}$+ | $-5.31 \times 10^{-7}$ |
| dijap | $-3.04 \times 10^{-7}$ | $-2.11 \times 10^{-7}$ |
| machiavelli | $7.41 \times 10^{-7}$* | $1.13 \times 10^{-6}$+ |

| | | |
|---|---|---|
| sagyfy | $5.60 \times 10^{-7}$* | $1.00 \times 10^{-7}$ |
| peeve | $-1.64 \times 10^{-7}$ | $-1.75 \times 10^{-7}$ |
| vegu | $-9.98 \times 10^{-7}$+ | $-2.24 \times 10^{-6}$* |
| DTC_ZM00013_consensus | $5.37 \times 10^{-7}$ | $-2.64 \times 10^{-6}$* |
| puck | $-9.40 \times 10^{-8}$ | $-1.31 \times 10^{-6}$* |
| DTC_ZM00101_consensus | $-4.44 \times 10^{-7}$ | $1.09 \times 10^{-8}$ |
| fourf | $6.44 \times 10^{-7}$ | $1.32 \times 10^{-6}$+ |
| DTC_ZM00085_consensus | $3.01 \times 10^{-7}$ | $1.19 \times 10^{-6}$ |
| guhis | $5.04 \times 10^{-8}$ | $1.45 \times 10^{-6}$* |
| Zm00743_AC177943_1 | $2.87 \times 10^{-7}$ | $-1.26 \times 10^{-6}$+ |
| ebel | $3.85 \times 10^{-10}$ | $-5.15 \times 10^{-7}$ |
| naseup | $5.45 \times 10^{-7}$ | $7.61 \times 10^{-7}$ |
| chr5_P_133655019 | $-9.49 \times 10^{-7}$* | $4.58 \times 10^{-7}$ |
| chr8_P_116327109 | $-4.96 \times 10^{-7}$ | $1.19 \times 10^{-7}$ |
| chr5_P_9712308 | $-2.56 \times 10^{-7}$ | $-1.82 \times 10^{-7}$ |
| dapuuv | $1.72 \times 10^{-6}$+ | $4.69 \times 10^{-6}$** |
| CRM1 | $-9.11 \times 10^{-7}$* | $1.47 \times 10^{-7}$ |
| iwik | $3.19 \times 10^{-6}$*** | $6.50 \times 10^{-6}$*** |
| DTC_ZM00038_consensus | $-3.14 \times 10^{-7}$ | $-1.53 \times 10^{-6}$ |
| chr2_D_48986571 | $3.86 \times 10^{-7}$ | $3.21 \times 10^{-7}$ |
| DTC_ZM00122_consensus | $-3.22 \times 10^{-8}$ | $-5.99 \times 10^{-7}$ |
| DTC_ZM00105_consensus | $1.03 \times 10^{-7}$ | $5.40 \times 10^{-6}$** |
| kameer | $3.02 \times 10^{-7}$ | $-1.23 \times 10^{-7}$ |
| DTC_ZM00012_consensus | $1.02 \times 10^{-6}$* | $1.15 \times 10^{-6}$ |
| neha | $-6.78 \times 10^{-7}$ | $-3.37 \times 10^{-7}$ |
| DTC_ZM00073_consensus | $-7.39 \times 10^{-7}$ | $-2.52 \times 10^{-6}$** |
| Zm00665_AC184764_1 | $2.56 \times 10^{-7}$ | $4.72 \times 10^{-7}$ |
| eninu | $-1.18 \times 10^{-6}$* | $-1.82 \times 10^{-6}$* |
| DTC_ZM00070_consensus | $-5.66 \times 10^{-7}$ | $2.64 \times 10^{-6}$+ |
| chr4_P_118808430 | $1.84 \times 10^{-7}$ | $1.69 \times 10^{-6}$* |
| DTC_ZM00102_consensus | $1.76 \times 10^{-6}$** | $2.33 \times 10^{-6}$+ |
| bipide | $-6.62 \times 10^{-7}$ | $-1.44 \times 10^{-6}$ |
| DTC_ZM00047_consensus | $-1.11 \times 10^{-6}$ | $-2.89 \times 10^{-6}$* |
| chr4_D_46804832 | $-3.10 \times 10^{-7}$ | $-8.71 \times 10^{-7}$ |
| DTC_ZM00053_consensus | $-4.60 \times 10^{-8}$ | $-2.04 \times 10^{-6}$ |
| chr1_P_193017007 | $-1.07 \times 10^{-6}$ | $7.94 \times 10^{-7}$ |
| ywyt | $6.93 \times 10^{-7}$ | $-3.51 \times 10^{-6}$** |
| lata | $8.89 \times 10^{-7}$ | $-4.49 \times 10^{-7}$ |
| wuge | $-3.27 \times 10^{-7}$ | $1.98 \times 10^{-6}$* |
| Hip1_19 | $1.70 \times 10^{-7}$ | $-5.01 \times 10^{-7}$ |
| DTC_ZM00091_consensus | $1.23 \times 10^{-7}$ | $1.50 \times 10^{-6}$ |
| Hip2_1 | $-9.21 \times 10^{-7}$ | $-2.62 \times 10^{-6}$+ |
| RIT_colonist_AC209460_0 | $2.58 \times 10^{-7}$ | $4.42 \times 10^{-6}$* |
| chr5_P_170822725 | $7.56 \times 10^{-7}$ | $1.41 \times 10^{-6}$ |
| TAFT1_DQ493649 | $1.48 \times 10^{-6}$ | $3.80 \times 10^{-8}$ |
| DTC_ZM00088_consensus | $-2.48 \times 10^{-8}$ | $-2.98 \times 10^{-7}$ |
| muojo | $-7.69 \times 10^{-7}$ | $-3.84 \times 10^{-6}$* |
| Zm00030_AC190592_1 | $1.44 \times 10^{-6}$** | $8.49 \times 10^{-7}$ |
| chr4_P_246400268 | $1.61 \times 10^{-7}$ | $-2.09 \times 10^{-6}$+ |
| anar | $1.36 \times 10^{-6}$+ | $2.46 \times 10^{-7}$ |
| DTH_ZM00024_consensus | $2.10 \times 10^{-6}$** | $-9.72 \times 10^{-7}$ |
| bosohe | $9.24 \times 10^{-7}$ | $4.88 \times 10^{-7}$ |
| CRM4 | $1.05 \times 10^{-6}$ | $-1.80 \times 10^{-7}$ |
| DTC_ZM00045_consensus | $-4.88 \times 10^{-7}$ | $2.80 \times 10^{-6}$* |
| DTC_ZM00044_consensus | $9.03 \times 10^{-7}$ | $3.62 \times 10^{-6}$ |
| uloh | $4.28 \times 10^{-7}$ | $-9.90 \times 10^{-7}$ |
| DTH_ZM00175_consensus | $-1.47 \times 10^{-6}$ | $-7.83 \times 10^{-6}$+ |

| | | |
|---|---|---|
| DTC_ZM00043_consensus | $7.40 \times 10^{-7}$ | $7.96 \times 10^{-7}$ |
| Hip2_2 | $-3.50 \times 10^{-7}$ | $2.86 \times 10^{-6}$+ |
| DTC_ZM00048_consensus | $-2.99 \times 10^{-7}$ | $-8.08 \times 10^{-7}$ |
| DTC_ZM00030_consensus | $2.87 \times 10^{-7}$ | $1.13 \times 10^{-6}$ |
| DTC_ZM00056_consensus | $5.53 \times 10^{-8}$ | $-4.33 \times 10^{-6}$*** |
| DTC_ZM00001_consensus | $-5.84 \times 10^{-7}$ | $1.54 \times 10^{-6}$ |
| bene | $1.85 \times 10^{-6}$ | $-3.51 \times 10^{-6}$ |
| Zm02117_AC177838_1 | $-2.30 \times 10^{-7}$ | $-2.33 \times 10^{-7}$ |
| Zm00268_AC191380_1 | $2.25 \times 10^{-6}$ | $9.32 \times 10^{-7}$ |
| dugiab | $-3.42 \times 10^{-6}$ | $-6.52 \times 10^{-6}$ |
| naiba | $-3.26 \times 10^{-6}$+ | $-6.62 \times 10^{-6}$+ |
| DTC_ZM00031_consensus | $-5.94 \times 10^{-7}$ | $-2.83 \times 10^{-6}$* |
| Zm02661_AC177816_1 | $-4.47 \times 10^{-7}$ | $-2.88 \times 10^{-6}$** |
| chr4_P_19323329 | $2.78 \times 10^{-7}$ | $5.06 \times 10^{-6}$*** |
| Zm02947_AC197071_1 | $7.11 \times 10^{-7}$ | $2.44 \times 10^{-6}$* |
| egisu | $-4.26 \times 10^{-7}$ | $-2.86 \times 10^{-6}$ |
| RIT_jare_AC204843_0 | $-8.71 \times 10^{-7}$ | $6.74 \times 10^{-7}$ |
| stonor | $-1.88 \times 10^{-6}$+ | $-2.83 \times 10^{-6}$ |
| Hip1_3 | $-5.10 \times 10^{-7}$ | $-5.45 \times 10^{-6}$* |
| DTC_ZM00002_consensus | $-8.43 \times 10^{-7}$ | $5.71 \times 10^{-7}$ |
| ibulaf | $-2.97 \times 10^{-8}$ | $-2.04 \times 10^{-6}$ |
| DTC_ZM00027_consensus | $5.53 \times 10^{-7}$ | $2.67 \times 10^{-6}$ |
| Zm00628_AC207419_1 | $-1.34 \times 10^{-7}$ | $-1.97 \times 10^{-6}$+ |
| DTH_ZM00035_consensus | $1.75 \times 10^{-6}$+ | $4.25 \times 10^{-6}$* |
| Zm02731_AC194595_1 | $-2.89 \times 10^{-7}$ | $8.07 \times 10^{-7}$ |
| RIL_otiot_AC206557_1 | $6.99 \times 10^{-7}$ | $-2.78 \times 10^{-6}$+ |
| Zm00364_AC196271_1 | $-3.94 \times 10^{-6}$* | $-1.79 \times 10^{-6}$ |
| DTC_ZM00026_consensus | $2.35 \times 10^{-6}$** | $7.54 \times 10^{-7}$ |
| pute | $4.75 \times 10^{-7}$ | $3.15 \times 10^{-6}$ |
| small | $-7.94 \times 10^{-7}$ | $-2.33 \times 10^{-6}$ |
| DTA_ZM00169_consensus | $1.62 \times 10^{-6}$ | $-7.15 \times 10^{-8}$ |
| DTA_ZM00135_consensus | $-2.91 \times 10^{-6}$ | $-1.12 \times 10^{-5}$* |
| DTA_ZM00060_consensus | $2.13 \times 10^{-6}$ | $-6.80 \times 10^{-6}$* |
| eugene | $1.89 \times 10^{-6}$ | $2.22 \times 10^{-6}$ |
| Zm07209_AC211743_1 | $-7.19 \times 10^{-7}$ | $6.78 \times 10^{-6}$+ |
| DTC_ZM00036_consensus | $-1.19 \times 10^{-7}$ | $2.08 \times 10^{-6}$ |
| DTC_ZM00051_consensus | $-2.42 \times 10^{-7}$ | $4.93 \times 10^{-7}$ |
| debeh | $-2.46 \times 10^{-7}$ | $-1.35 \times 10^{-6}$ |
| DTH_ZM00004_consensus | $-1.98 \times 10^{-7}$ | $5.52 \times 10^{-6}$** |
| Zm00473_AC191070_1 | $-3.07 \times 10^{-6}$ | $-1.13 \times 10^{-5}$** |
| ewib | $4.69 \times 10^{-6}$** | $1.13 \times 10^{-6}$ |
| DTA_ZM00373_consensus | $-3.64 \times 10^{-6}$ | $4.93 \times 10^{-6}$ |
| liove | $-2.05 \times 10^{-7}$ | $-4.86 \times 10^{-6}$** |
| RIX_ekeje_AC197537_0 | $-2.91 \times 10^{-7}$ | $-3.34 \times 10^{-6}$ |
| DTC_ZM00003_consensus | $-1.90 \times 10^{-6}$+ | $-5.45 \times 10^{-6}$** |
| CRM3 | $1.66 \times 10^{-6}$ | $3.45 \times 10^{-6}$+ |
| DTH_ZM00021_consensus | $4.67 \times 10^{-6}$ | $7.08 \times 10^{-7}$ |
| RIL_nubo_AC212389_0 | $8.81 \times 10^{-7}$ | $2.93 \times 10^{-6}$ |
| RIL_afeda_AC208414_0 | $-1.27 \times 10^{-6}$ | $4.00 \times 10^{-6}$ |
| kake | $1.50 \times 10^{-6}$ | $2.80 \times 10^{-6}$ |
| hutu | $-1.72 \times 10^{-6}$ | $-4.86 \times 10^{-7}$ |
| Hip1_26 | $-9.01 \times 10^{-7}$ | $-7.00 \times 10^{-7}$ |
| DTA_ZM00383_consensus | $-3.26 \times 10^{-6}$ | $-2.05 \times 10^{-6}$ |
| name | $4.35 \times 10^{-6}$ | $1.60 \times 10^{-5}$* |
| bogu | $7.40 \times 10^{-7}$ | $5.73 \times 10^{-6}$** |
| DTH_ZM00102_consensus | $-2.30 \times 10^{-6}$ | $7.80 \times 10^{-6}$ |
| DTC_ZM00100_consensus | $3.13 \times 10^{-7}$ | $-1.26 \times 10^{-6}$ |

| | | |
|---|---|---|
| DTC_ZM00063_consensus | $-4.40 \times 10^{-6}$* | $-8.63 \times 10^{-6}$* |
| DTH_ZM00032_consensus | $-2.78 \times 10^{-6}$* | $-1.14 \times 10^{-6}$ |
| DTC_ZM00034_consensus | $-2.10 \times 10^{-6}$ | $6.32 \times 10^{-7}$ |
| DTA_ZM00291_consensus | $-4.31 \times 10^{-6}$ | $-1.03 \times 10^{-5}$* |
| Num.Obs. | 1454 | 1561 |
| R2 | 0.265 | 0.540 |
| R2 Adj. | 0.166 | 0.483 |

+ $p < 0.1$, * $p < 0.05$, ** $p < 0.01$, *** $p < 0.001$

**Table S8.** Effect sizes of the relationship between TE base pairs of each of the 170 TE families with greater than 10 Mb of sequence across all NAM individuals, and flowering time (DTS) or grain yield (GY).
